# 15-deoxy-Δ^12,14^-prostaglandin J_2_ limits *Salmonella* infection through regulation of host TLR4 signaling and inflammasome activation

**DOI:** 10.64898/2026.08.24.746850

**Authors:** Nathalia Santos Magalhães, Viktoriia V. Feofanova, Vivian T. Nguyen, Heidi Pauer, Larissa D. Silva Ferreira, Gabriela Ceccon Chianca, L. Caetano M. Antunes

## Abstract

Enteric infections caused by *Salmonella enterica* remain a major global health concern and are increasingly associated with antimicrobial resistance. Therefore, new strategies to combat this important pathogen are needed. The interactions between *S. enterica* and the human host have been the subject of intense investigation over the last several decades, yet new findings continue to emerge. We previously showed that 15-deoxy-Δ^12,14^-prostaglandin J_2_ (15d-PGJ_2_) reduces *Salmonella* colonization of macrophages, but the mechanisms underlying this protective effect were still unknown. Here, we demonstrate that 15d-PGJ_2_ limits *Salmonella* infection by suppressing TLR4 signaling and inflammasome activation. Treatment with 15d-PGJ_2_ reduced TLR4 expression, NF-κB activation, iNOS, COX-2, nitric oxide production, IL-1β release, and inflammasome-related targets, including NLRP3 and caspase-1 activity, while only partially reversing macrophage polarization. Combined treatment with the TLR4 antagonist TAK-242 further reduced bacterial colonization of and IL-1β release by macrophages, supporting the involvement of TLR4 signaling in the effects of 15d-PGJ_2_. During mouse infections, 15d-PGJ_2_ reduced bacterial burdens in a tissue-dependent manner. Together, these findings demonstrate that 15d-PGJ_2_ limits *Salmonella* infection through selective modulation of TLR4 signaling and inflammasome activation.

## Introduction

Enteric infections are a major public health concern, particularly due to the increasing antimicrobial resistance displayed by enteric pathogens (1). One of the most concerning infectious diarrheal diseases worldwide is salmonellosis, caused by *Salmonella enterica*, a Gram-negative, rod-shaped, facultative intracellular bacterium. Several serovars of *S. enterica* are currently included in the list of major pathogens for antimicrobial resistance in global public health, including *S. enterica* serovar Typhimurium (2). In the United States, it is estimated that approximately 1.35 million people are infected each year, resulting in about 420 deaths (3).

The main cellular targets for infection and replication of *Salmonella* are innate immune cells, particularly macrophages. *Salmonella* is recognized by receptors expressed on macrophages through the binding of pathogen-associated molecular patterns (PAMPs) expressed by the bacterium, such as lipopolysaccharide (LPS) and flagellin. One of the most extensively studied receptors involved in this interaction is Toll-like receptor 4 (TLR4) (4). In general, during bacterial infection, bacterial cells are recognized by macrophages and internalized into a phagosome, which subsequently fuses with lysosomes, leading to pathogen killing and elimination. However, due to bacterial escape mechanisms, especially encoded by the type III secretion system (T3SS) within the *Salmonella* Pathogenicity Island 2 (SPI-2), *Salmonella* is able to survive within the *Salmonella*-containing vacuole (SCV) by preventing phagosome maturation (5). In addition, *Salmonella* can modulate the host immune response to promote its replication and spread to new cells (6, 7). One example of this modulation is the alteration of reactive oxygen and nitrogen species production, as well as cytokine production by the host cell (8–12). Among the cytokines most upregulated during the course of infection are interleukins (IL)-1, IL-6, IL-12, and IL-18, as well as tumor necrosis factor (TNF)-α (13).

In a previous study from our group, we showed that 15-deoxy-Δ^12,14^-prostaglandin J_2_ (15d-PGJ_2_) reduced the colonization of macrophages by *Salmonella*, although the underlying mechanism was not elucidated (14). 15d-PGJ_2_ is a lipid mediator derived from arachidonic acid through the activity of the cyclooxygenase-2 (COX-2) enzyme. Although its anti-inflammatory effects have been extensively described in cancer and metabolic diseases (15–18), the role of this bioactive lipid in the context of bacterial infection remains unclear. 15d-PGJ_2_ is recognized as an agonist of peroxisome proliferator-activated receptor gamma (PPARγ), which partially explains its anti-inflammatory properties. However, it has also been demonstrated that this prostaglandin can exert biological effects independently of PPARγ. In our previous work, we showed that co-treatment of macrophages with a PPARγ inhibitor and 15d-PGJ_2_ did not reverse the reduction in *Salmonella* colonization induced by 15d-PGJ_2_, suggesting that this effect occurs independently of PPARγ (14).

Beyond its role as a PPARγ agonist, 15d-PGJ_2_ has emerged as an important modulator of innate immune signaling pathways. Consistent with this, reports in the literature show that 15d-PGJ_2_ reduces the expression and activation of TLR4 through the inhibition of nuclear factor (NF)-κB in pancreatic acinar cells in a model of acute pancreatitis (19). TLR4 activation has been recognized as a key upstream event in inflammasome activation during both infectious and non-infectious inflammatory responses (20, 21). The inflammasome is a multiprotein complex activated in response to inflammatory stimuli that promotes caspase activation, leading to the maturation of the pro-inflammatory cytokines IL-1β and IL-18. The canonical inflammasome complex is composed of a nucleotide-binding domain and a leucine-rich repeat-containing receptor (NLR), such as NLRP3 or NLRC4, the adaptor protein ASC, and the effector protease caspase-1. Upon assembly of this complex, caspase-1 cleaves pro-IL-1β and pro-IL-18 into their active forms, which are subsequently released by the cells. Given its central role in coordinating innate immune responses, the TLR4-inflammasome signaling axis has been extensively investigated in bacterial infections, including those caused by *Salmonella*.

In murine *Salmonella* infection models, TLR4/MyD88-dependent innate signaling and inflammasome activation contribute to intestinal inflammation and host defense. *Salmonella* activates caspase-1 and promotes IL-1β and IL-18 production through NLRC4 and NLRP3 inflammasomes, and intestinal upregulation of NLRP3 and IL-1β has also been documented (22–24). Similarly, Guan *et al*. demonstrated that *Salmonella* infection in poultry led to increased expression of key components of the TLR4 signaling pathway, including TLR4, MyD88, and NF-κB. Concomitantly, infection was associated with upregulation of intestinal inflammasome-related markers, such as NLRP3, caspase-1, IL-1β, and IL-18 (25). Moreover, it has been observed that 15d-PGJ_2_ decreases inflammasome pathway activity in macrophages and in a murine model challenged with anthrax lethal toxin by reducing caspase-1 activation and IL-1β release (26). Together, these observations suggest that modulation of the TLR4-inflammasome axis may contribute to the protective effects of 15d-PGJ_2_ during *Salmonella* infection.

*Salmonella* is recognized as a potent activator of the inflammasome pathway (27). Although the role of inflammasome activation in the resolution of infection remains controversial, accumulating evidence suggests that *Salmonella* can exploit this pathway as a mechanism of immune evasion (28). Inflammasome activation is also closely associated with pyroptosis, a lytic form of programmed cell death. Upon activation, caspase-1 cleaves gasdermin D (GSDMD), which is responsible for pore formation in the plasma membrane. These pores facilitate the release of mature IL-1β and other intracellular contents. However, when this response is excessive or dysregulated, GSDMD-mediated pore formation can lead to uncontrolled cell lysis and the massive release of intracellular components, further amplifying the inflammatory response. In this context, *Salmonella* may take advantage of inflammasome-induced pyroptosis to promote its dissemination to neighboring cells, thereby sustaining and propagating the infection (29).

Collectively, these previous findings led us to hypothesize that modulation of TLR4 signaling and downstream inflammasome activation contributes to the protective effects of 15d-PGJ_2_ during *Salmonella* infection. To test this hypothesis, we investigated the impact of 15d-PGJ_2_ on TLR4 signaling, inflammasome activation, and bacterial colonization in macrophages and in a murine model of infection.

## Material and methods

### Bacterial and eukaryotic cell culture conditions

All experiments used *Salmonella* enterica serovar Typhimurium strain SL1344. To start *Salmonella* cultures, frozen stocks stored at −80°C were used to streak Luria-Bertani (LB; Fisher BioReagents, Waltham, MA, USA) agar plates containing 100 μg/mL streptomycin (SL1344 is intrinsically resistant to this antibiotic). Plates were incubated overnight at 37°C. Subsequently, a single colony was inoculated into LB broth containing streptomycin and incubated at 37°C with shaking (200 rpm).

Macrophages (RAW 264.7) were purchased from the American Type Culture Collection (ATCC, Manassas, VA, USA). HEK-Dual™ cells (NF-κB and ISG Dual Reporter HEK 293 Cells) were obtained from InvivoGen (San Diego, CA, USA). RAW 267.4 and HEK 293 cells were maintained in Dulbecco’s Modified Eagle Medium (DMEM; Thermo Fisher Scientific, Waltham, MA, USA) supplemented with 10% (v/v) heat-inactivated fetal bovine serum (FBS; Cytiva, Marlborough, MA, USA) and 1% (v/v) penicillin (100 U/mL) and streptomycin (100 μg/mL) (Thermo Fisher Scientific), at 37°C with 5% CO_2_. Cells were maintained up to a maximum of 20 passages.

### MTT and lactate dehydrogenase assays

RAW 267.4 cells were seeded in 96-well plates in DMEM supplemented with 10% FBS and incubated at 37°C with 5% CO_2_. The following day, the medium was removed, and 200 µL of medium containing 15d-PGJ_2_ was added, in the presence or absence of 10% FBS, as indicated, and cells were then returned to the incubator. For analysis of cellular metabolic activity, 22 hours after the addition of 15d-PGJ_2_, 5 µg/mL 3-(4,5-dimethylthiazol-2-yl)-2,5-diphenyltetrazolium bromide (MTT; Sigma-Aldrich, St. Louis, MO, USA) was added to the cells and incubated for 2 hours. At the end of the 24-hour experiment, the plate was centrifuged at 300 × g for 5 minutes at 4°C, and the supernatant was discarded. Finally, the cells were resuspended in dimethyl sulfoxide (DMSO), and absorbance was measured by spectrophotometry at 570 nm using a SpectraMax i3 spectrophotometer (Molecular Devices, San Jose, CA, USA). The results were normalized to the control group mean. For cytotoxicity analysis, CytoTox 96® kit (Promega, Madison, WI, USA) was used to measure the amount of lactate dehydrogenase (LDH) present in supernatants. Because intact cells do not release LDH, the quantification of this enzyme in the supernatant is directly proportional to cell lysis. The same cell seeding protocol, 15d-PGJ_2_ doses, and experimental time points performed for the MTT assay were used for LDH quantification. All manufacturers’ instructions were followed for each assay.

### Bacterial growth curves

For the analysis of *Salmonella* growth curves, 1 µL of an overnight culture of strain SL1344 was added to 200 µL of LB broth containing the appropriate antibiotic in a 96-well plate in the presence or absence of 15d-PGJ_2_. The plate was incubated at 37°C in a spectrophotometric plate reader for 24 hours (SpectraMax i3 spectrophotometer; Molecular Devices), and the optical density at 600 nm (OD_600_) was measured every 30 minutes.

### Tissue culture infections

RAW 267.4 cells or HEK-Dual™ cells were seeded in plates containing DMEM supplemented with 10% FBS and incubated overnight at 37°C with 5% CO_2_. On the following day, an overnight *Salmonella* culture was measured for optical density (OD_600_). The bacterial culture was then centrifuged at 3200 × g for 3 minutes. The supernatant was discarded, and the bacterial pellet was washed with 1X PBS and centrifuged as mentioned previously. Subsequently, *Salmonella* was resuspended in 1 mL of DMEM at the appropriate density. Host cells were infected at a multiplicity of infection (MOI) of 10. After 30 minutes, the cells were washed with 1X PBS and incubated with 100 μg/mL gentamicin for 1 hour and 30 minutes. After, the medium was replaced with a lower gentamicin concentration (10 μg/mL) and maintained for a total of 24 hours of infection. For colony-forming unit (CFU) assays, cells were seeded in 96-well plates at a density of 3 × 10 cells per well in DMEM supplemented with 10% FBS. The cells were infected the following day as described above. After 24 hours post-infection, the supernatant was collected, and a 0.05% Triton solution in 1X PBS was added to lyse the eukaryotic cells and expose the bacteria present in the intracellular compartment. From this lysate, serial dilutions were prepared and plated on LB agar plates. Plates were incubated overnight at 37°C, and the number of CFU was determined.

For tissue culture experiments, cells were treated with a range of doses of 15d-PGJ_2_ (2, 5, or 10 µM; Cayman Chemical, Ann Arbor, MI, USA) for 24 hours, depending on the experiment. The H-PGDS inhibitor and T0070907 (both from Cayman Chemical) were used at a final concentration of 1 µM and remained present throughout the 24-h infection period. The concentration of IL-1β (Thermo Fisher Scientific) used was 1, 5, or 10 ng/mL, and it remained present throughout the 24 hours of infection. The concentration of Nigericin (InvivoGen) used in the cells was 10 µM. For this treatment, nigericin was added 1 hour before infection and maintained throughout the entire experiment, resulting in a total of 25 hours of stimulation. For experiments using TAK-242 (1 µM; Cayman Chemical), the following strategy was applied: cells were seeded in the morning, and at the end of the day they were treated overnight with TAK-242 to block TLR4. On the following day, the medium was removed and the infection protocol was performed, maintaining the TAK-242 treatment throughout the entire experiment.

### Animal experiments

C57BL/6 mice (6-8 weeks old) were purchased from The Jackson Laboratory (Bar Harbor, ME, USA). Upon arrival, mice were acclimated for a minimum of three days in the University of Kansas Animal Care Unit. Animals were housed in groups of up to five in a temperature-controlled (22 °C ± 2 °C), humidity- and light-controlled (12-h light/dark period) colony room, with *ad libitum* access to food and water. Animals were randomly divided into four groups: uninfected (n = 5), uninfected + 15d-PGJ_2_ (n = 5), infected (n = 5), and infected + 15d-PGJ_2_ (n = 5). On day 1 of the experiment, a high dose of streptomycin (20 mg in 100 μL) was administered by oral gavage in all animals to disrupt the gut microbiota. On the following day, mice were infected with approximately 10^8^ CFU of *Salmonella* (strain SL1344). For this, *Salmonella* was grown overnight at 37°C with shaking in LB broth containing streptomycin (100 μg/mL). Bacterial cells were then centrifuged and resuspended in PBS to a density of 10^9^ cells/mL, and 100 μL was administered to each animal by oral gavage. On the same day of infection, animals were treated with vehicle (saline) or 15d-PGJ_2_ (1 mg/kg, intraperitoneally). Mice were treated daily throughout the duration of the experiment. After four days of infection, mice were humanely euthanized. Spleen, liver, intestines, and feces were collected, weighed, and processed or stored. For CFU analyses, tissue samples were placed into tubes containing beads and 1 mL of PBS. Organs were homogenized using a TissueLyser III (QIAGEN, Hilden, Germany). Homogenates were serially diluted in PBS and plated on LB agar containing streptomycin (100 μg/mL). Plates were incubated overnight at 37°C, and the number of CFU was counted and normalized by tissue weight.

This protocol was approved by the Institutional Animal Care and Use Committee (IACUC) of the University of Kansas (protocol number: 2403001671).

### NF-κB activation reporter

For NF-κB activation analysis, HEK-Dual™ cells were seeded in 96-well plates and incubated overnight at 37°C with 5% CO_2_. On the following day, cells were infected at an MOI of 10, as previously described. At the end of the 24-hour infection period, cell supernatants were collected, the QUANTI-Blue™ Solution (InvivoGen, San Diego, CA, USA) was added, and the mixture was incubated for 2 hours at 37°C. Optical density (OD) was then measured at 630 nm using a microplate reader (SpectraMax i3 spectrophotometer; Molecular Devices).

### Nitrite oxide quantification

For reactive nitric oxide (NO) analysis, we measured levels of nitrite, a stable metabolite of NO, in the supernatant of cells using the Griess Reagent System Kit (Promega). All the manufacturers’ recommendations were followed.

### Quantitative Real-Time PCR (qRT-PCR)

For experiments with RAW 264.7 cells, cells were seeded in 6-well plates and incubated at 37°C with 5% CO_2_ overnight. The following day, cells were infected at an MOI of 10, as previously described. After 24 hours of infection, cells were washed with PBS and RNA was extracted using the RNeasy Mini Kit (QIAGEN). For tissue samples, organs were stored in RNAprotect Tissue Reagent (QIAGEN) at −80°C until processing. Tissues were removed from RNAprotect and transferred to tubes containing lysis buffer and beads, and dissociation was performed using a TissueLyser III (QIAGEN). Subsequently, RNA extraction was carried out using the RNeasy Mini Kit (QIAGEN) according to the manufacturer’s instructions, including the optional DNase treatment step (RNase-Free DNase Set, QIAGEN) to remove genomic DNA contamination. To evaluate the direct impact of 15d-PGJ_2_ on bacterial gene expression, *Salmonella* SL1344 was grown overnight in LB medium at 37°C under shaking conditions (200 rpm). The culture was then diluted 1:200 in fresh LB broth with or without 10 µM 15d-PGJ_2_ and incubated at 37°C with shaking (200 rpm) for 4 hours. Following incubation, bacterial RNA was stabilized using RNAprotect Bacteria Reagent (QIAGEN) according to the manufacturer’s instructions. Bacterial cells were subsequently pelleted by centrifugation, and total RNA was extracted using the RNeasy Mini Kit (QIAGEN) following the manufacturer’s recommendations. To remove residual genomic DNA, RNA samples were treated with Turbo DNase (2 U/µL; Life Technologies, Waltham, MA, USA), followed by an additional purification step using the same RNeasy Mini Kit (QIAGEN), according to the manufacturer’s instructions. From this point onward, all subsequent procedures were performed identically for RNA samples obtained from RAW 264.7 cells, tissue specimens, and bacterial cultures, and followed the manufacturer’s recommendations. RNA concentrations were determined using an ND-1000 NanoDrop spectrophotometer (Thermo Fisher Scientific). cDNA synthesis was performed using the GoScript™ Reverse Transcription System (Promega). Quantitative real-time PCR (qRT-PCR) was conducted using the PowerTrack™ SYBR™ Green Master Mix (Applied Biosystems, Waltham, MA, USA) on a QuantStudio™ 3 Real-Time PCR System (Applied Biosystems). To confirm the absence of genomic DNA contamination, cDNA synthesis reactions without reverse transcriptase were performed in parallel, and the absence of amplification was verified during qRT-PCR. Gene expression was normalized to the housekeeping genes GAPDH for host cells and *gapA* for bacterial samples. Relative expression levels were calculated using the 2^-ΔΔCt^ method (30). Primer sequences are listed in Table S1.

### Cytokine quantification

TNF-α, G-CSF, CCL22, CXCL1, CCL17, IL-6, IL-12p70, IL-12p40, IL-18, IL-23, and free active TGF-β1 levels were measured in cell supernatants using the LEGENDplex™ Mouse Macrophage/Microglia Panel (BioLegend, San Diego, CA, USA). All procedures were performed according to the manufacturer’s instructions. For IL-1β quantification by ELISA, an IL-1 beta/IL-1F2 Elisa Kit (R&D Systems, Minneapolis, MN, USA) was used and all procedures were performed according to the manufacturer’s instructions.

For cytokine analysis in cell culture samples, supernatants were collected and stored at −20°C until used. For animal cecum samples, tissues were stored at −80°C until the day of processing. On the day of processing, RIPA buffer supplemented with phosphatase and protease inhibitors was added to the tissues in the presence of beads, and tissues were lysed using the TissueLyser III. Two cycles of 5 minutes were performed to promote tissue dissociation. Samples were centrifuged at 10,000 × g for 10 minutes at 4°C, and the supernatant was collected and stored at −80°C. Cytokine levels were normalized by tissue weight.

### Statistical analysis

Results were presented as mean ± standard error of the mean (SEM). Data were first assessed for normal distribution and then analyzed using one-way analysis of variance (ANOVA), followed by the Tukey multiple comparisons post hoc test. When only two groups were compared, unpaired Student’s t-tests were used. Statistical analyses were performed using GraphPad Prism 10 software. A *p*-value of ≤ 0.05 was considered statistically significant.

## Results

### 15d-PGJ_2_ acts in the later stages of Salmonella infection in cultured macrophages

We previously showed that the addition of exogenous 15d-PGJ_2_ to macrophages infected with *Salmonella* caused a reduction in bacterial counts at several time points up until 24 hours of infection (14). Here, we set out to first confirm these original findings while also extending our infection times to determine the timing of the effects of 15d-PGJ_2_ on bacterial burden. By doing so, we confirmed our original report that the addition of 15d-PGJ_2_ affected *Salmonella* during macrophage infection by showing a dose dependent reduction in bacterial counts. The maximum effect of 15d-PGJ_2_ on bacterial colonization in *Salmonella*-infected macrophages was observed at 10 µM (Fig. 1A). It is important to note that this concentration did not induce any cytotoxicity in macrophages (Fig. S1A and S1B), even under reduced FBS supplementation or in the complete absence of FBS (Fig. S1C). Additionally, we observed that the effect of 15d-PGJ_2_ was more pronounced at 24 hours post-infection, with a reduction of 75% in bacterial counts (Fig. 1B). Conversely, 15d-PGJ_2_ caused a reduction in *Salmonella* burdens of 28 and 55% at 36 and 48 hours post-infection, respectively, compared to no significant effect or a slight increase in bacterial loads at 2, 4, 6, 8, and 12 hours post-infection. This suggested that the production of 15d-PGJ_2_ is somehow detrimental to *Salmonella* survival and/or replication inside macrophages. To confirm this, we used a chemical inhibitor of the Hematopoietic Prostaglandin D Synthase (H-PGDS), the enzyme responsible for the synthesis of 15d-PGJ_2_, and showed that treatment of macrophages with this inhibitor significantly increased bacterial loads during infection, supporting a protective role for this pathway against *Salmonella*. Importantly, 15d-PGJ_2_ did not affect *Salmonella* growth *in vitro* (Fig. S2A and S2B), indicating that its effect on colonization is host-dependent. Together, these results indicate that 15d-PGJ_2_ limits *Salmonella* colonization at later stages of infection.

**Figure 1.**
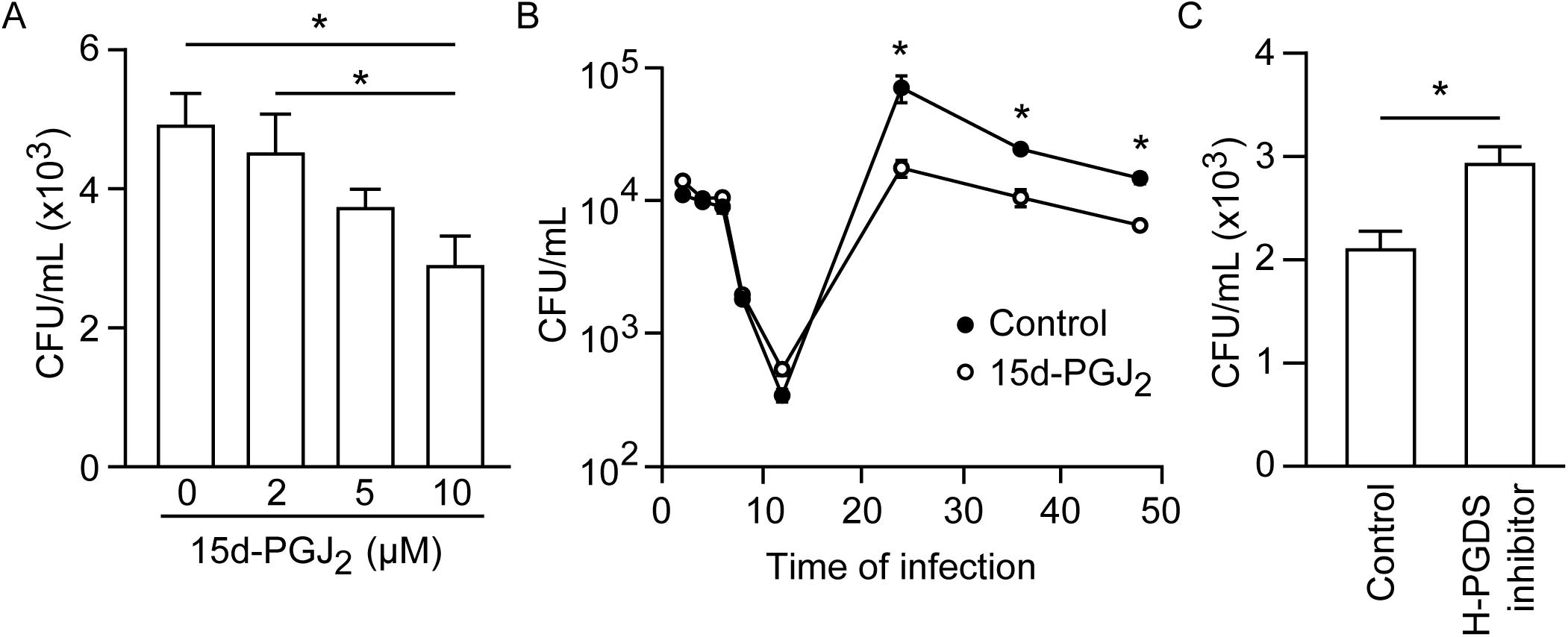
15d-PGJ_2_ decreases *Salmonella* colonization in macrophages. (A) *Salmonella* colonization in macrophages was assessed by quantification of colony-forming units (CFU) at different concentrations of 15d-PGJ_2_ (2, 5, and 10 µM) 24 hours after infection. (B) The impact of 15d-PGJ_2_ (10 µM) on *Salmonella* burdens was evaluated at different time points after infection. (C) The effect of a H-PGDS inhibitor (1 µM) on *Salmonella* burdens was evaluated 24 hours after infection. An MOI of 10 was used in all experiments described. The data shown are from one representative experiment out of three independent experiments performed under the same conditions. Each independent experiment included 5 replicates per condition. Bars represent the means ± standard errors of the means (SEM) of the 5 replicates from the representative experiment shown. Statistical analyses were performed using Student’s t-test for comparisons between two groups and one-way ANOVA followed by Tukey’s multiple comparisons test for comparisons involving three or more groups. ns, non-significant; \**p*<0.05. h.p.i., hours post-infection; H-PGDS, hematopoietic prostaglandin D synthase.

### 15d-PGJ_2_ reverts the proinflammatory profile of macrophages during Salmonella infection

We next investigated the effect of *Salmonella* infection on the expression of several inflammatory mediators in macrophages, using qRT-PCR. *Salmonella* infection induced the expression of iNOS (Fig. 2A), COX-2 (Fig. 2C), IL-23 (Fig. 2E), IL-10 (Fig. 2G), IL-1β (Fig. 2H), NLRP3 (Fig. 2J), MyD88 (Fig. 2Q), and p65 (Fig. 2R) in macrophages 24 hours after infection compared to the uninfected group. In contrast, the expression of Arg1 (Fig. 2B), IL-6 (Fig. 2F), ASC (Fig. 2K), and GSDMD (Fig. 2M) was repressed by infection under the same conditions. No significant differences were observed in the expression of PPARγ (Fig. 2D), TNF-α (Fig. 2I), caspase-1 (Fig. 2L), IL-18 (Fig. 2N), TLR2 (Fig. 2O), TLR4 (Fig. 2P), and NF-κB (Fig. 2S). These results indicate that *Salmonella* infection modulates inflammatory and inflammasome-related pathways in macrophages.

**Figure 2.**
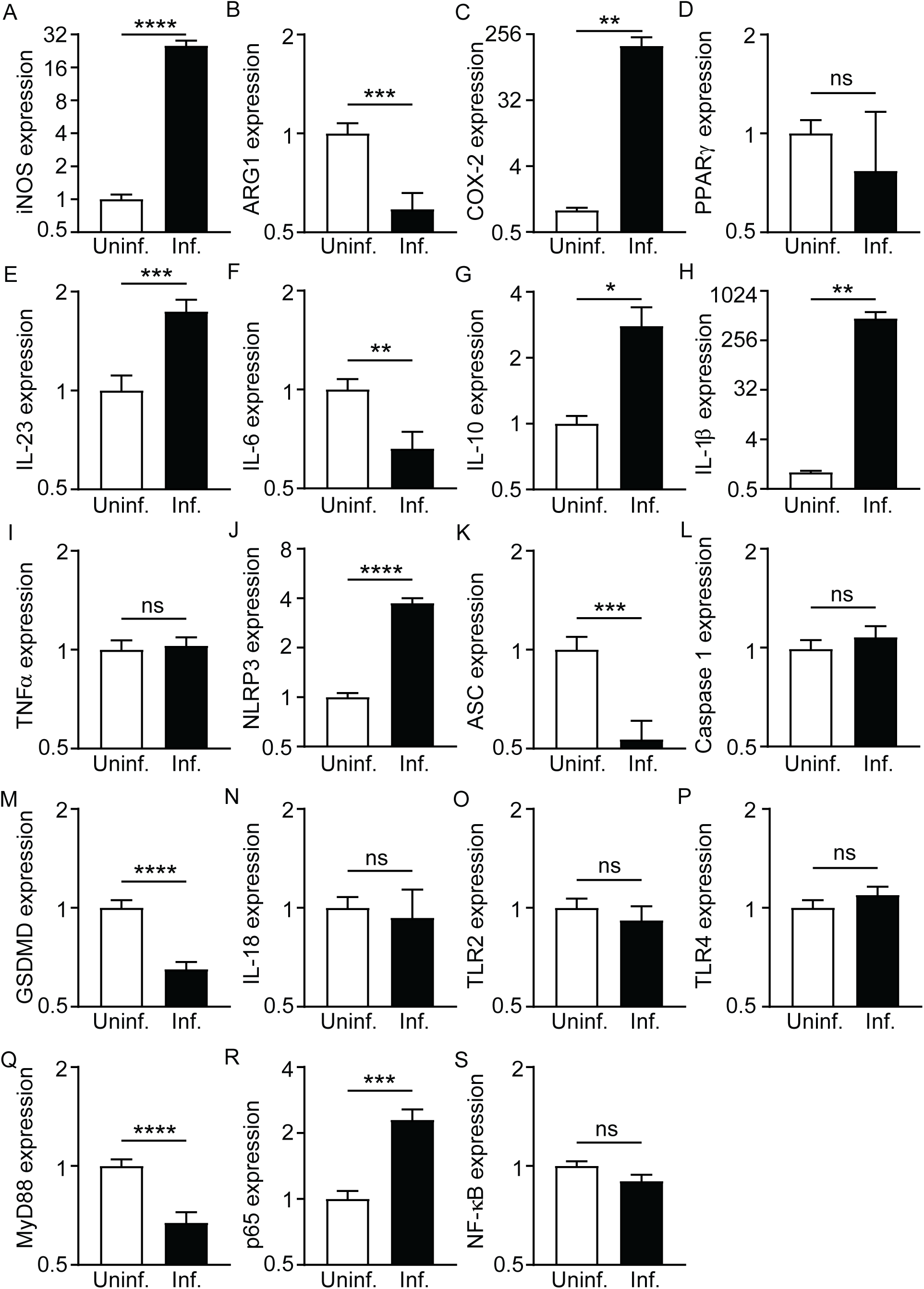
Salmonella modulates important inflammatory genes during macrophage infection. Expression levels of (A) iNOS, (B) Arg1, (C) COX-2, (D) PPARγ, (E) IL-23, (F) IL-6, (G) IL-10, (H) IL-1β, (I) TNF-α, (J) NLRP3, (K) ASC, (L) caspase-1, (M) GSDMD, (N) IL-18, (O) TLR2, (P) TLR4, (Q) MyD88, (R) p65, and (S) NF-κB were analyzed by qRT-PCR 24 hours after infection. An MOI of 10 was used. Data were normalized to the housekeeping gene GAPDH and presented as log_2_ fold change relative to the uninfected group. Data shown are pooled from three independent experiments, each performed with 5 replicates per condition. Bars represent the means ± standard errors of the means (SEM) of the pooled data. Statistical analyses were performed using Student’s t-test. ns, non-significant; \**p*<0.05; \*\**p*<0.01; \*\*\**p*<0.001; \*\*\*\**p*<0.0001. iNOS, inducible nitric oxide synthase; Arg1, arginase 1; COX-2, cyclooxygenase-2; PPARγ, peroxisome proliferator-activated receptor gamma; IL, interleukin; TNF-α, tumor necrosis factor alpha; NLRP3, NOD-like receptor family pyrin domain-containing 3; ASC, apoptosis-associated speck-like protein containing a CARD; GSDMD, gasdermin D; TLR, Toll-like receptor; MyD88, myeloid differentiation primary response 88; NF-κB, nuclear factor kappa-light-chain-enhancer of activated B cells.

To determine whether 15d-PGJ_2_ modulates the inflammatory response induced by *Salmonella*, we then analyzed the expression of key markers of macrophage activation during infection, in the absence or presence of 15d-PGJ_2_ supplementation. As mentioned above, *Salmonella*-infected, untreated cells showed increased expression of iNOS and COX-2, and 15d-PGJ_2_ treatment significantly reduced their expression (Fig. 3A and 3C). Consistent with this, nitric oxide (NO) production was elevated in infected cells, but reduced to basal levels upon treatment with 15d-PGJ_2_ (Fig. 3E). In contrast, the repression of anti-inflammatory markers such as Arg1 (Fig. 3B) and PPARγ (Fig. 3D) by infection was not restored by 15d-PGJ_2_ treatment. In addition, treatment with a PPARγ antagonist did not reverse the effects of 15d-PGJ_2_ on bacterial colonization (Fig. S3A) or NO production (Fig. S3B), reinforcing the notion that 15d-PGJ_2_ acts through a PPARγ-independent mechanism to modulate *Salmonella* infection. Altogether, these findings indicate that 15d-PGJ_2_ does not fully reverse macrophage polarization but instead selectively modulates important inflammatory pathways. At the cytokine level, transcriptional and secreted responses showed distinct patterns. *Salmonella* infection decreased IL-6 (Fig. 4A) and increased IL-23 (Fig. 4B) expression, and these responses were not significantly altered by 15d-PGJ_2_. However, no differences in IL-6 (Fig. S4E) or IL-23 (Fig. S4I) secretion were observed between groups. Similarly, no changes were detected in the release of G-CSF, CCL22, CXCL1, CCL17, IL-12p70, IL-12p40, IL-18, or free active TGF-β1 (Fig. S4A-J). In contrast, IL-1β expression (Fig. 4C) and secretion (Fig. 4F) were increased upon infection and significantly reduced by 15d-PGJ_2_ treatment. Finally, IL-10 and TNF-α regulation also showed divergence between transcriptional and protein levels. Although *Salmonella* infection caused no significant changes at the mRNA level (Fig. 4D and 4E), IL-10 (Fig. 4G) and TNF-α (Fig. 4H) secretion was significantly increased during infection, an effect that was significantly reduced by 15d-PGJ_2_ treatment. Together, these results indicate that 15d-PGJ_2_ promotes a selective reprogramming of macrophage inflammatory responses rather than a complete reversal of their activation state.

**Figure 3.**
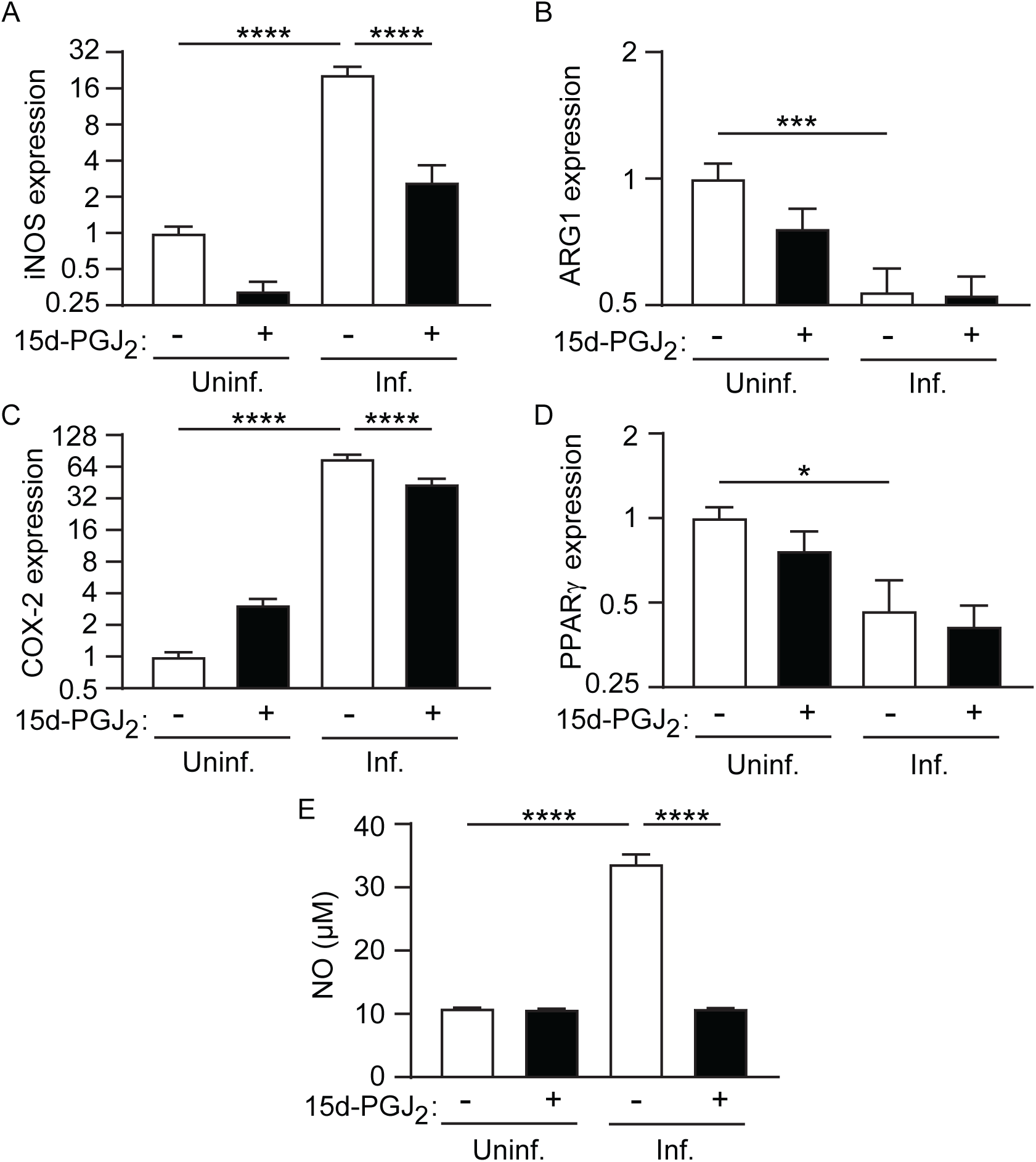
15d-PGJ_2_ dampens the proinflammatory profile of *Salmonella*-infected macrophages. The effect of 15d-PGJ_2_ (10 µM) on (A) iNOS, (B) Arg1, (C) COX-2, and (D) PPARγ expression was analyzed by qRT-PCR 24 hours after infection. Data were normalized to the housekeeping gene GAPDH and presented as log_2_ fold change relative to the uninfected group. (E) Nitrate quantification was performed using the Griess Reagent System Kit. An MOI of 10 was used. Data shown are pooled from two independent experiments, each performed with 5 replicates per condition. Bars represent the means ± standard errors of the means (SEM) of the pooled data. Statistical analyses were performed using one-way ANOVA followed by Tukey’s multiple comparisons test. ns, non-significant; \**p*<0.05; \*\*\**p*<0.001; \*\*\*\**p*<0.0001. iNOS, inducible nitric oxide synthase; Arg1, arginase 1; COX-2, cyclooxygenase-2; PPARγ, peroxisome proliferator-activated receptor gamma; NO, nitric oxide.

**Figure 4.**
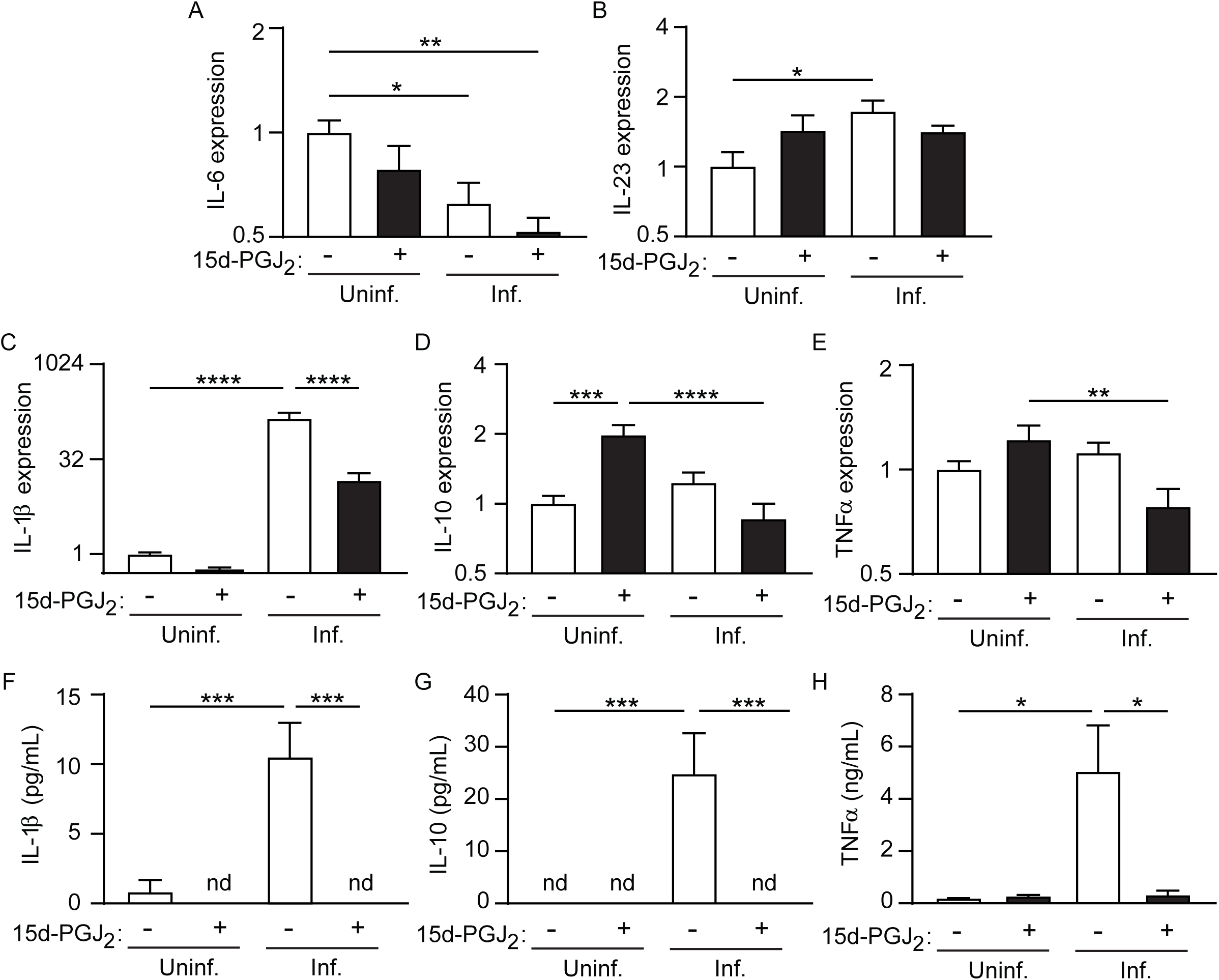
15d-PGJ_2_ downregulates cytokine production in *Salmonella*-infected macrophages. The effect of 15d-PGJ_2_ (10 µM) on (A) IL-6, (B) IL-23, (C) IL-1β, (D) IL-10, and (E) TNF-α expression was analyzed by qRT-PCR 24 hours after infection. Data were normalized to the housekeeping gene GAPDH and presented as log_2_ fold change relative to the uninfected group. (F) IL-1β and (G) IL-10 were quantified by ELISA 24 hours after infection. Data shown in panels A-G are pooled from two independent experiments, each performed with 5 replicates per condition. (H) TNF-α levels were quantified using the LEGENDplex™ Mouse Macrophage/Microglia Panel Kit 24 hours after infection. Data shown in panel H are from a single experiment performed with four replicates per condition. An MOI of 10 was used. Bars represent the means ± standard errors of the means (SEM); for panels A-G, means ± SEM were calculated from the pooled data. Statistical analyses were performed using one-way ANOVA followed by Tukey’s multiple comparison test. ns, non-significant; \**p*<0.05; \*\**p*<0.01; \*\*\**p*<0.001; \*\*\*\**p*<0.0001. IL, interleukin; TNF-α, tumor necrosis factor alpha; n.d., not detected.

### 15d-PGJ_2_ regulates the macrophage inflammasome pathway during Salmonella infection

The production and secretion of IL-1β is a hallmark of inflammasome activation (31). Given that IL-1β was strongly modulated by 15d-PGJ_2_, we next investigated the impact of this eicosanoid on other components of the inflammasome pathway. *Salmonella* infection increased NLRP3 expression (Fig. 5A), and this effect was partially reduced by 15d-PGJ_2_. In contrast, ASC (Fig. 5B) and GSDMD (Fig. 5D) were downregulated in infected cells, with further reduction of GSDMD upon 15d-PGJ_2_ treatment. Additionally, 15d-PGJ_2_ decreased the expression of ASC and GSDMD in uninfected macrophages. Caspase-1 showed a divergence between transcription and activity data. While mRNA levels were unchanged (Fig. 5C), active caspase-1 was increased in infected cells (Fig. 5F), indicating post-transcriptional regulation of inflammasome activation. As for IL-1β, 15d-PGJ_2_ treatment reverted the activation of caspase-1 in infected cells. Several *Salmonella* genes have been shown to be involved in the modulation of inflammasome activation by the pathogen (27, 32). Therefore, we tested whether 15d-PGJ_2_ can affect the expression of some of these bacterial genes. Our data showed that none of the *Salmonella* genes associated with inflammasome modulation showed differences in expression during bacterial growth in the absence or presence of 15d-PGJ_2_ (Fig. S5A-R), supporting a mechanism of action that is solely driven by the host response.

**Figure 5.**
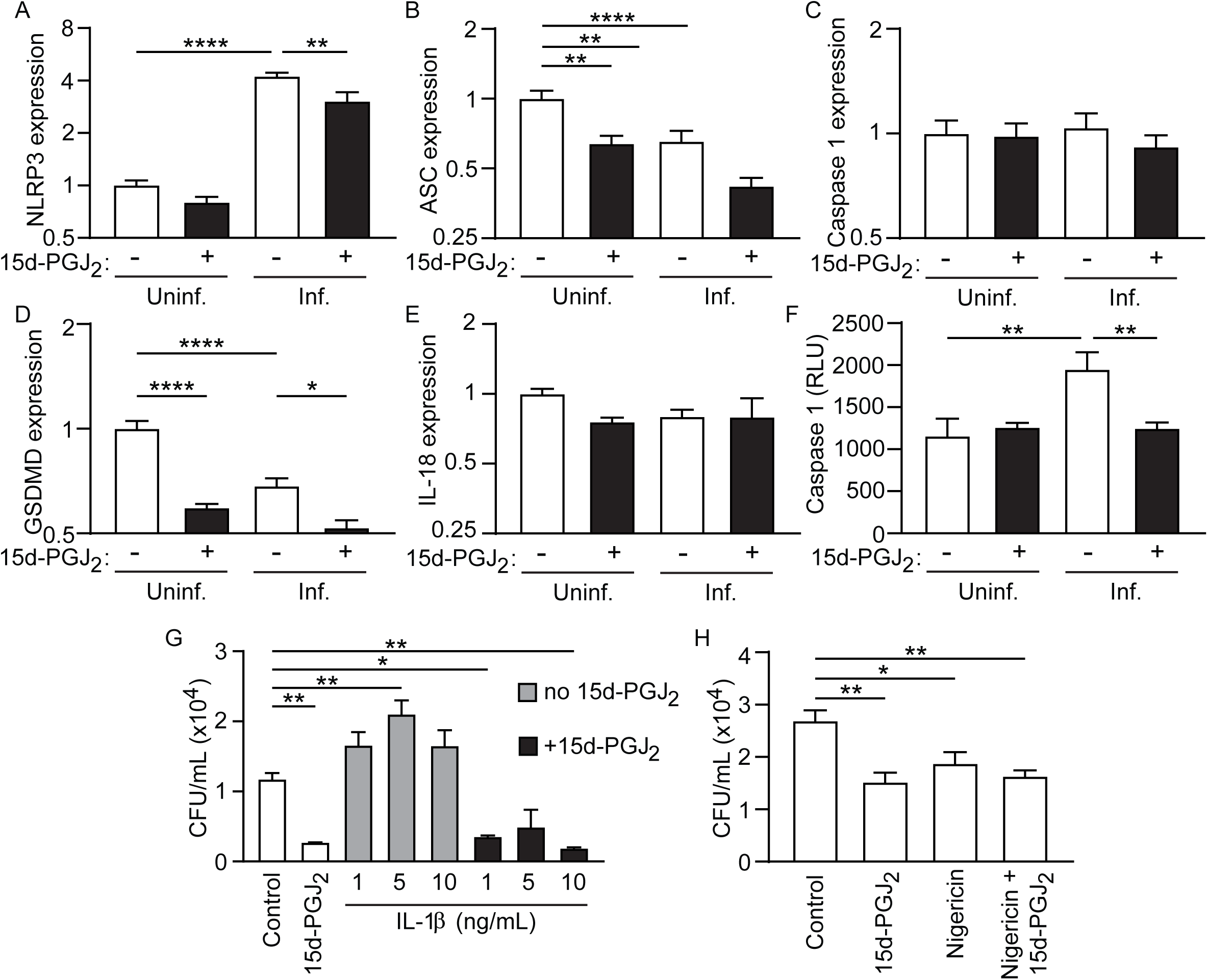
15d-PGJ_2_ downregulates the inflammasome pathway in *Salmonella*-infected macrophages. The effect of 15d-PGJ_2_ (10 µM) on (A) NLRP3, (B) ASC, (C) caspase-1, (D) GSDMD, and (E) IL-18 expression was analyzed by qRT-PCR 24 hours after infection. Data were normalized to the housekeeping gene GAPDH and presented as log_2_ fold change relative to the uninfected group. (F) Active caspase-1 was quantified by bioluminescence using the Caspase-Glo® 1 Inflammasome Assay. Data shown in panels A-F are pooled from two independent experiments, each performed with 5 replicates per condition. Bars represent the means ± standard errors of the means (SEM) of the pooled data. The effects of (G) IL-1β and (H) nigericin on *Salmonella* colonization were evaluated by CFU quantification 24 hours after infection. Panels G and H show data from one representative experiment out of three independent experiments performed under the same conditions. Each independent experiment included 5 replicates per condition. Bars represent the means ± SEM of the 5 replicates from the representative experiment shown. An MOI of 10 was used. Statistical analyses were performed using one-way ANOVA followed by Tukey’s multiple comparisons test. ns, non-significant; \**p*<0.05; \*\**p*<0.01; \*\*\*\**p*<0.0001. NLRP3, NOD-like receptor family pyrin domain-containing 3; ASC, apoptosis-associated speck-like protein containing a CARD; GSDMD, gasdermin D; IL, interleukin; CFU, colony-forming units.

Given that bacterial burdens correlated with IL-1β levels in our experiments, we next set out to test if exogenous IL-1β supplementation could rescue *Salmonella* colonization in 15d-PGJ_2_-treated macrophages. Our data showed that, although IL-1β treatment caused a significant increase in bacterial colonization in the absence of 15d-PGJ_2_, the addition of up to 10 ng/mL of recombinant IL-1β to infected macrophages did not cause a significant change in bacterial burdens in the presence of 15d-PGJ_2_ (Fig. 5G). This prompted us to investigate if inflammasome activation *in trans* with nigericin would affect *Salmonella* burdens in 15d-PGJ_2_-treated macrophages. As with IL-1β treatment, nigericin-induced inflammasome activation did not counteract the reduction in colonization caused by 15d-PGJ_2_ (Fig. 5H). Together, these findings indicate that although 15d-PGJ_2_ modulates inflammasome activation during infection, its effect on *Salmonella* colonization likely involves other pathways.

### 15d-PGJ_2_ regulates the crosstalk between the TLR4 and inflammasome pathways during Salmonella infection

During *Salmonella* infection, TLR4 signaling is known to contribute to inflammatory activation, including inflammasome responses. To investigate whether 15d-PGJ_2_ modulates this pathway, we first evaluated TLR expression. Although TLR4 mRNA levels were not increased upon infection (Fig. 6B), 15d-PGJ_2_ treatment significantly reduced TLR4 expression in infected macrophages compared to untreated *Salmonella*-infected cells. A similar pattern was observed for TLR2 expression (Fig. 6A). We next analyzed downstream components of the TLR4 signaling pathway. *Salmonella* infection downregulated MyD88 (Fig. 6C) and upregulated p65 (Fig. 6D) expression compared to uninfected controls. However, 15d-PGJ_2_ treatment reduced the expression of both targets in infected macrophages. Similarly, NF-κB mRNA levels were not altered by infection but were significantly reduced upon 15d-PGJ_2_ treatment (Fig. 6E). To functionally assess NF-κB activation, we used an NF-κB reporter cell line (HEK-Dual™). *Salmonella* infection increased NF-κB activation, whereas 15d-PGJ_2_ treatment partially reversed this effect (Fig. 6F). Consistent with previous findings, 15d-PGJ_2_ also reduced bacterial colonization (Fig. 6G). However, the reduced activation of NF-κB was not solely attributable to reduced *Salmonella* burden, as NF-κB activation remained significantly lower following 15d-PGJ_2_ treatment even after normalization by bacterial load (Fig. 6H). Likewise, when transcriptional data were normalized by bacterial load, iNOS and IL-1β expression remained significantly reduced following 15d-PGJ_2_ treatment (Fig. S6A-T), indicating that these effects are independent of differences in bacterial burden. To further investigate the role of TLR4 signaling, we evaluated the effect of a TLR4 antagonist on *Salmonella* infection. TLR4 blockade alone did not significantly reduce *Salmonella* colonization compared to the control (DMSO). However, the combination of the TLR4 antagonist with 15d-PGJ_2_ further decreased bacterial load compared to 15d-PGJ_2_ treatment alone (Fig. 7A). In parallel, this combined treatment further reduced IL-1β release compared to cells treated only with 15d-PGJ_2_ (Fig. 7B). However, the combination of TLR4 antagonist and 15d-PGJ_2_ did not further reduce caspase-1 activation in infected cells compared to 15d-PGJ_2_ treatment alone (Fig. 7C). Together, these results suggest that 15d-PGJ_2_ modulates the TLR4-NF-κB axis and its crosstalk with the inflammasome pathway. The enhanced reduction in bacterial colonization observed with the treatment of 15d-PGJ_2_-treated, *Salmonella*-infected cells with a TLR4 antagonist suggests that 15d-PGJ_2_ and the TLR4 antagonist may act through partially independent mechanisms, supporting the role of 15d-PGJ_2_ as a selective immunomodulator rather than a single-pathway inhibitor.

**Figure 6.**
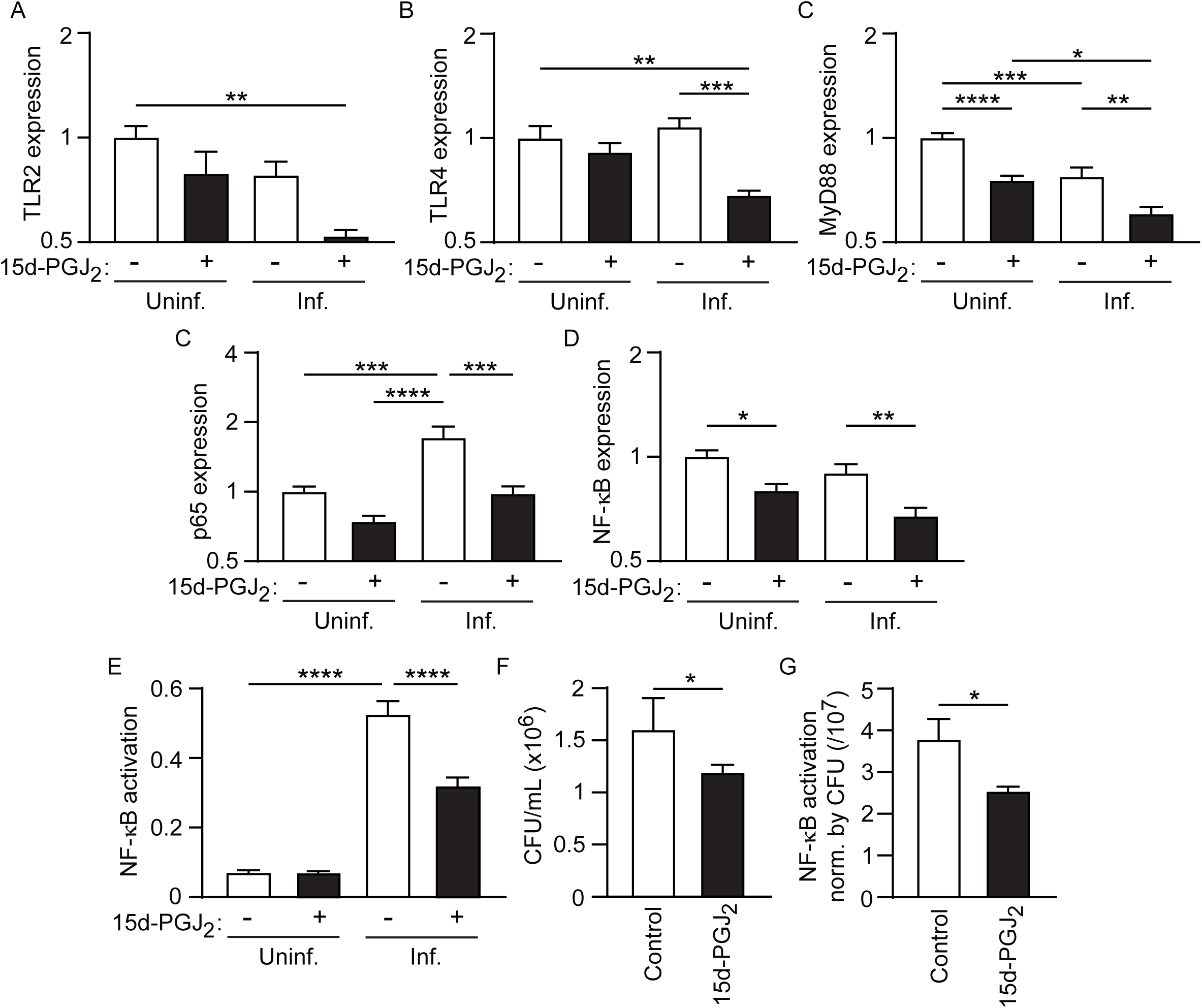
15d-PGJ_2_ downregulates the TLR4 pathway in *Salmonella*-infected cells. The effect of 15d-PGJ_2_ (10 µM) on (A) TLR2, (B) TLR4, (C) MyD88, (D) p65, and (E) NF-κB expression was analyzed by qRT-PCR 24 hours after infection. Data were normalized to the housekeeping gene GAPDH and presented as log_2_ fold change relative to the uninfected group. Data shown in panels A-E are pooled from two independent experiments, each performed with 5 replicates per condition. Bars represent the means ± standard errors of the means (SEM) of the pooled data. (F) NF-κB activation was quantified using NF-κB reporter cells. (G) The impact of 15d-PGJ_2_ on *Salmonella* colonization in NF-κB reporter cells was evaluated by CFU quantification. (H) The ratios of NF-κB activation to CFUs observed was determined in NF-κB reporter cells 24 hours after infection. Panels F-H show data from one representative experiment out of three independent experiments performed under the same conditions. Each independent experiment included 5 replicates per condition. Bars represent the means ± SEM of the 5 replicates from the representative experiment shown. In panel G, outliers were identified and removed using the ROUT method (Q = 1%) prior to statistical analysis. An MOI of 10 was used. Statistical analyses were performed using one-way ANOVA followed by Tukey’s multiple comparisons test. ns, non-significant; \**p*<0.05; \*\**p*<0.01; \*\*\**p*<0.001; \*\*\*\**p*<0.0001. TLR, Toll-like receptor; MyD88, myeloid differentiation primary response 88; NF-κB, nuclear factor kappa-light-chain-enhancer of activated B cells; CFU, colony-forming units.

**Figure 7.**
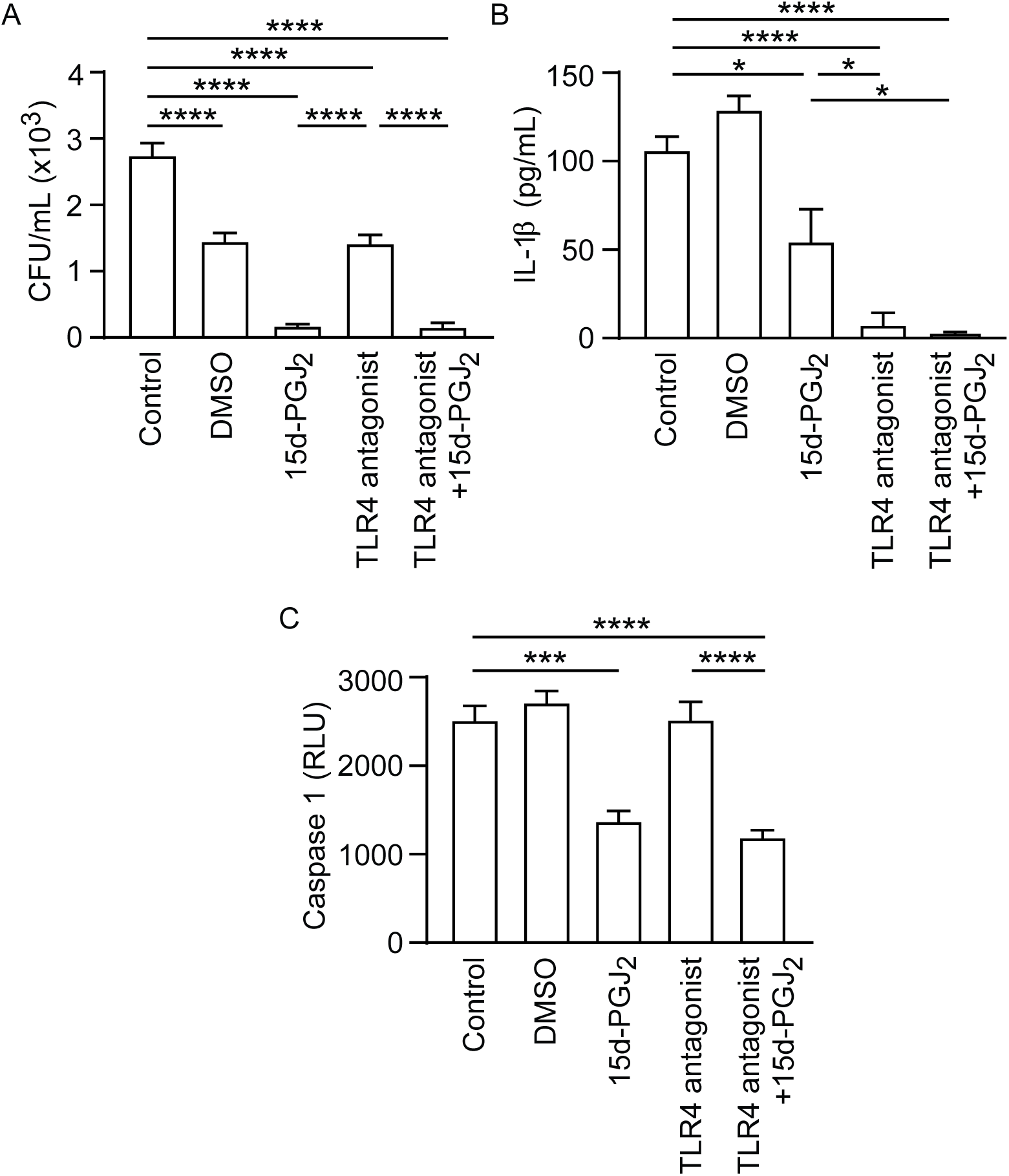
15d-PGJ_2_ downregulates the crosstalk between the TLR4 and inflammasome pathways in *Salmonella*-infected macrophages. (A) The impact of the combination of 15d-PGJ_2_ and the TLR4 antagonist TAK-242 (1 µM) on bacterial colonization and (B) IL-1β production in *Salmonella*-infected macrophages was analyzed 24 hours after infection by CFU quantification and ELISA, respectively. (C) Active caspase-1 was quantified by bioluminescence using the Caspase-Glo® 1 Inflammasome Assay. As a control group for TAK-242, cells were treated with the same amount of vehicle (DMSO). An MOI of 10 was used. The graphs show data from one representative experiment out of two independent experiments performed under the same conditions. Each independent experiment included 5 replicates per condition. Bars represent the means ± standard errors of the means (SEM) of the five replicates from the representative experiment shown. Statistical analyses were performed using the Student’s t-test for comparisons between two groups and one-way ANOVA followed by Tukey’s multiple comparisons test for comparisons involving three or more groups. ns, non-significant; \**p*<0.05; \*\**p*<0.01; \*\*\**p*<0.001; \*\*\*\**p*<0.0001. TLR, Toll-like receptor; IL, interleukin; CFU, colony-forming units; DMSO, dimethyl sulfoxide.

### Endogenous 15d-PGJ_2_ contributes to the control of Salmonella colonization in macrophages

To investigate whether endogenous 15d-PGJ_2_ production contributes to protection against *Salmonella* colonization, we treated macrophages with an H-PGDS inhibitor in the presence or absence of exogenous 15d-PGJ_2_. As expected, treatment with 15d-PGJ_2_ alone significantly decreased bacterial loads, whereas inhibition of endogenous 15d-PGJ_2_ production increased *Salmonella* colonization in macrophages. Notably, the combination of the H-PGDS inhibitor with exogenous 15d-PGJ_2_ maintained bacterial burden at levels comparable to the 15d-PGJ_2_-treated group, suggesting that exogenous 15d-PGJ_2_ can compensate for the inhibition of its endogenous production (Fig. 8A). We next evaluated IL-1β release under these conditions. Although increased bacterial loads were observed in macrophages treated with the H-PGDS inhibitor, no significant differences in IL-1β release were detected compared to untreated cells. In contrast, treatment with 15d-PGJ_2_, either alone or in combination with the H-PGDS inhibitor, reduced IL-1β levels to below the detection limit of the assay (Fig. 8B). Together, these results suggest that endogenous 15d-PGJ_2_ contributes to the control of *Salmonella* colonization, while exogenous 15d-PGJ_2_ is sufficient to override the effects of its pharmacological inhibition.

**Figure 8.**
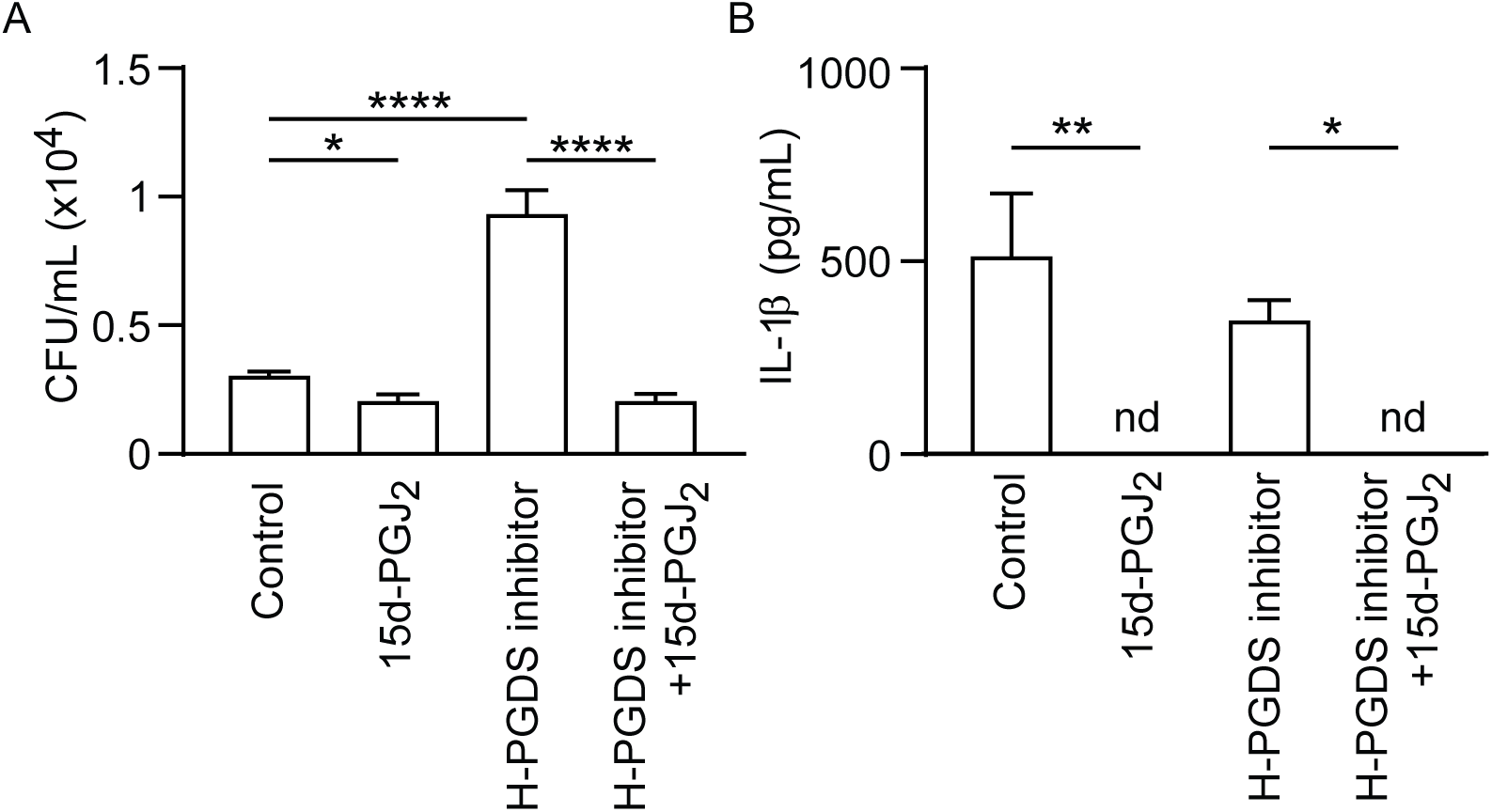
Blockade of endogenous 15d-PGJ_2_ production enhances bacterial colonization but does not affect IL-1β release in *Salmonella*-infected macrophages. (A) Impact of the H-PGDS inhibitor (1 µM) on macrophage colonization by *Salmonella* was evaluated by CFU quantification 24 hours after infection. (B) IL-1β production was measured in infected cell supernatants by ELISA 24 hours after infection. An MOI of 10 was used. The graphs show data from one representative experiment out of two independent experiments performed under the same conditions. Each independent experiment included 5 replicates per condition. Bars represent the means ± standard errors of the means (SEM) of the five replicates from the representative experiment shown. Statistical analyses were performed using one-way ANOVA followed by Tukey’s multiple comparisons test. \**p*<0.05; \*\**p*<0.01; \*\*\*\**p*<0.0001. H-PGDS, hematopoietic prostaglandin D synthase; CFU, colony-forming units; IL, interleukin; n.d., not detected.

### 15d-PGJ_2_ decreases Salmonella colonization of the murine cecum

To investigate the role of 15d-PGJ_2_ during *Salmonella* infection in a complex *in vivo* model, we performed infection experiments in mice, as illustrated in Fig. 9A. *Salmonella*-infected mice showed reduced body weight (Fig. S7A) and decreased lengths of the cecum, colon, and large intestine (Fig. S7C and S7D) compared to the untreated control group, as expected. However, 15d-PGJ_2_ treatment did not reverse these parameters. Representative images of the large intestine are shown in Fig. S7B. We next evaluated bacterial colonization in different organs 4 days post-infection. *Salmonella* burden was assessed in the spleen (Fig. 9B), liver (Fig. 9C), cecum (Fig. 9D), colon (Fig. 9E), and feces (Fig. 9F). Notably, 15d-PGJ_2_ treatment selectively reduced bacterial loads in the cecum, with no significant effects observed in the other tissues analyzed. Together, these results suggest that 15d-PGJ_2_ reduces *Salmonella* colonization *in vivo*, with a localized effect in the cecum.

**Figure 9.**
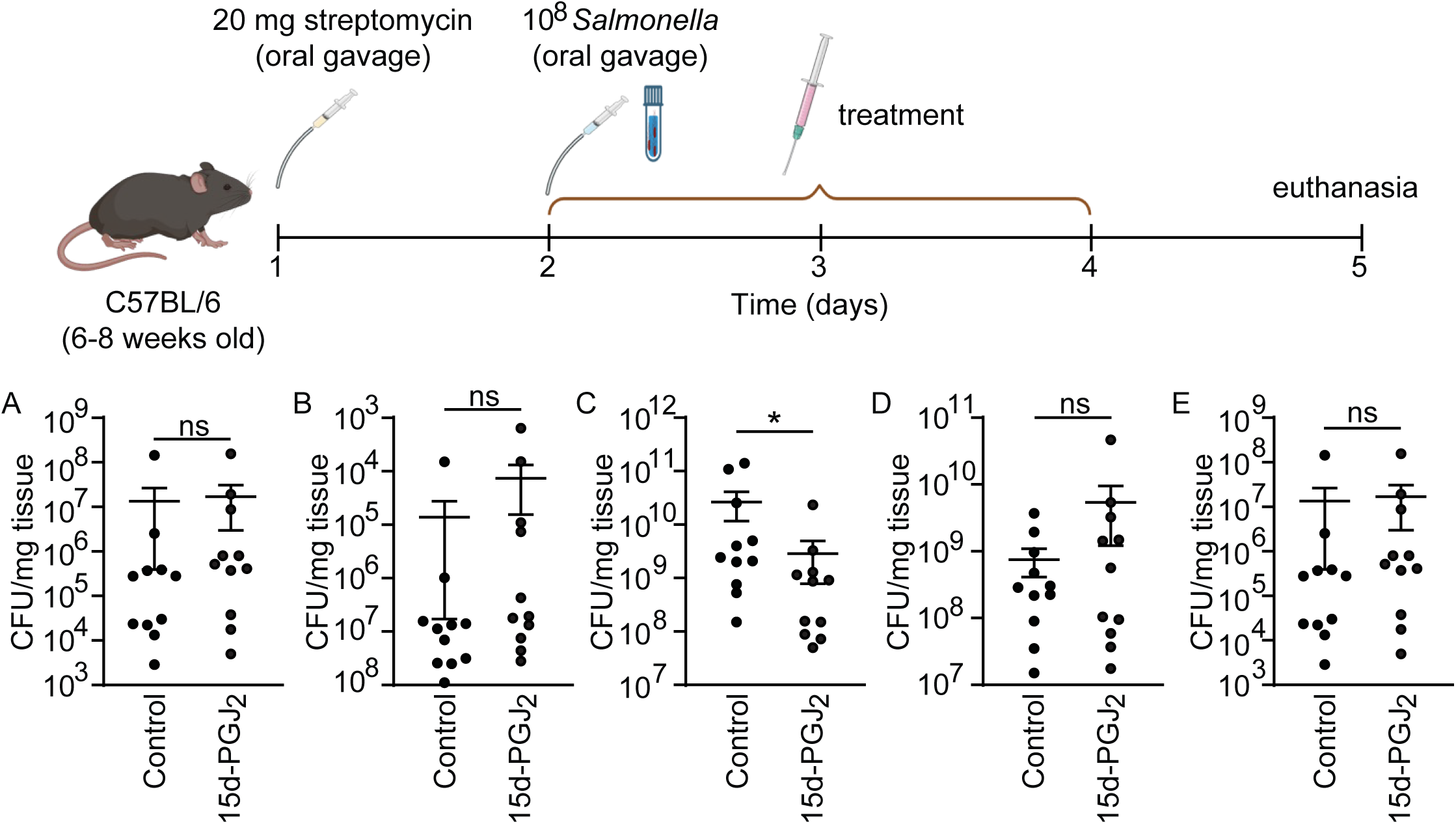
15d-PGJ_2_ treatment decreases bacterial loads in the cecum of *Salmonella*-infected mice. (A) The cartoon illustrates a schematic flowchart of the mouse experiment. The graphs represent the impact of 15d-PGJ_2_ (1 mg/kg) treatment on bacterial colonization in the spleen (B), liver (C), cecum (D), colon (E), and feces (F) 4 days post-infection. Data were normalized to the weight of the corresponding organ. Each dot represents an individual animal. Data shown are pooled from two independent experiments, with 5 or 6 mice per experiment. Bars represent the means ± standard errors of the means (SEM). Statistical analyses were performed using the Student’s t-test. \**p*<0.05. CFU, colony-forming units.

### In vivo inflammatory profiles suggest context-dependent regulation of 15d-PGJ_2_ during Salmonella infection

To investigate whether the inflammatory targets modulated by 15d-PGJ_2_ *in vitro* were also regulated *in vivo*, we analyzed the expression levels of several immune genes in the ceca of *Salmonella*-infected mice. Expression levels of iNOS, Arg1, COX-2, PPARγ, IL-23, IL-6, IL-10, IL-1β, TNF-α, NLRP3, ASC, Caspase-1, GSDMD, IL-18, TLR2, TLR4, MyD88, p65, and NF-κB (Fig. 10A-T) were evaluated 4 days post-infection by qRT-PCR. Interestingly, oral 15d-PGJ_2_ treatment increased the expression of COX-2 (Fig. 10C), TNF-α (Fig. 10I), NLRP3 (Fig. 10J), TLR2 (Fig. 10O), TLR4 (Fig. 10P), MyD88 (Fig. 10R), and NF-κB (Fig. 10T) in *Salmonella*-infected mice compared to untreated controls. In contrast, GSDMD expression was decreased in the 15d-PGJ_2_-treated group (Fig. 10N). Together, these results indicate that, although 15d-PGJ_2_ reduces *Salmonella* colonization *in vivo*, its modulation of inflammatory gene expression in the cecum differs from that observed *in vitro*, suggesting a context-dependent regulation of host responses.

**Figure 10.**
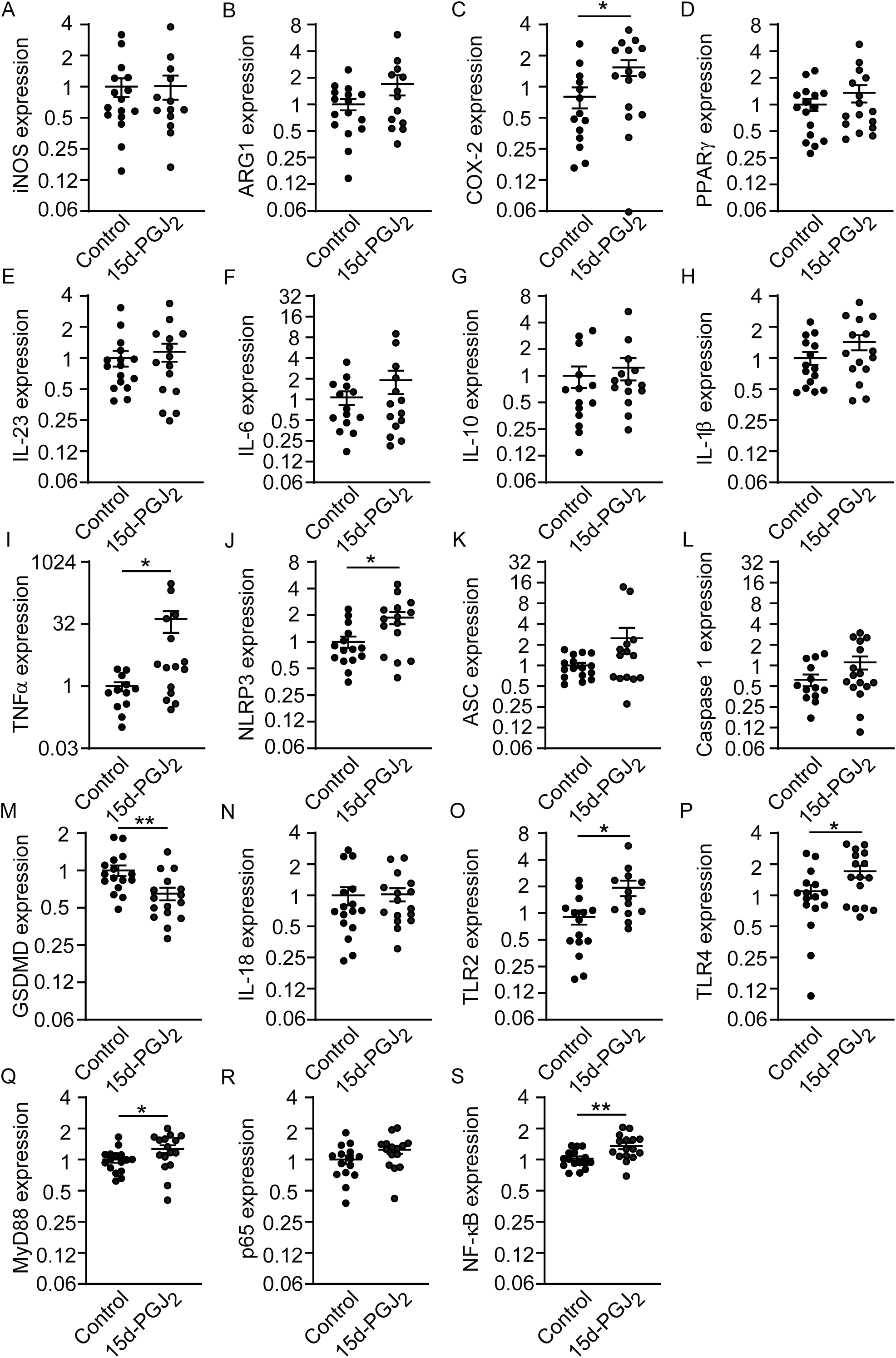
15d-PGJ_2_ treatment modulates key inflammatory genes in the cecum of *Salmonella*-infected mice. The effect of 15d-PGJ_2_ (1 mg/kg) on (A) iNOS, (B) Arg1, (C) COX-2, (D) PPARγ, (E) IL-23, (F) IL-6, (G) IL-10, (H) IL-1β, (I) TNF-α, (J) NLRP3, (K) ASC, (L) caspase-1, (M) GSDMD, (N) IL-18, (O) TLR2, (P) TLR4, (Q) MyD88, (R) p65, and (S) NF-κB expression in the cecum was analyzed by qRT-PCR 4 days post-infection. Data were normalized to the housekeeping gene GAPDH and presented as log_2_ fold change relative to the infected group. Each dot represents an individual animal. Data shown are pooled from three independent experiments, with 5 or 6 mice per experiment. Outliers were identified and removed using the ROUT method (Q = 1%) prior to statistical analysis. Statistical analyses were performed using the Student’s t-test. ns, non-significant; \**p*<0.05; \*\**p*<0.01. iNOS, inducible nitric oxide synthase; Arg1, arginase 1; COX-2, cyclooxygenase-2; PPARγ, peroxisome proliferator-activated receptor gamma; IL, interleukin; TNF-α, tumor necrosis factor alpha; NLRP3, NOD-like receptor family pyrin domain-containing 3; ASC, apoptosis-associated speck-like protein containing a CARD; GSDMD, gasdermin D; TLR, Toll-like receptor; MyD88, myeloid differentiation primary response 88; NF-κB, nuclear factor kappa-light-chain-enhancer of activated B cells.

## Discussion

15d-PGJ_2_ is an anti-inflammatory lipid mediator whose roles are well studied in cancer and chronic diseases. However, its role in infectious diseases is still poorly understood. Previously, our group showed that 15d-PGJ_2_ reduces *Salmonella* colonization in macrophages, although the underlying mechanisms remained unclear (14). In this study, we show that 15d-PGJ_2_ reduces *Salmonella* colonization in both *in vitro* and *in vivo* models, and that this effect appears to involve modulation of the TLR4-inflammasome pathway.

Previous data from our group showed that *Salmonella* infection increases endogenous 15d-PGJ_2_ levels in cultured macrophages (14) as well as during mouse infections (33). Here, we show that inhibition of the H-PGDS enzyme further increases bacterial burdens in macrophages, suggesting that *Salmonella* colonization is at least in part controlled by endogenous production of 15d-PGJ_2_. Additionally, *Salmonella* infection has also been shown to increase COX-2 expression, promoting the production of several eicosanoids, including 15d-PGJ_2_. Yang *et al.* demonstrated that inhibition of H-PGDS plays distinct roles in the clearance of *Escherichia coli* from infected macrophages depending on the time point analyzed. At early stages of infection, H-PGDS inhibition reduced bacterial colonization and was associated with decreased NF-κB activation; however, the opposite effect was observed at later stages (34). These findings suggest that components of this pathway may shift their role in immune modulation over the course of infection.

15d-PGJ_2_ is well known as an activator of PPARγ; however, the data presented in this study support and expand our previous observation that the effects of 15d-PGJ_2_ on *Salmonella* macrophage infection are PPARγ-independent (14). Also, our results show that 15d-PGJ_2_ decreases bacterial loads while modulating inflammatory responses in macrophages, including iNOS and COX-2 expression, nitric oxide (NO) production, cytokine expression, and NF-κB activation. Consistent with this, previous studies have shown that 15d-PGJ_2_ inhibits both NF-κB and AP-1 activity at the iNOS promoter, which may partially explain our findings (35). Besides, 15d-PGJ_2_ was previously implicated in the attenuated iNOS expression and IL-1β production in *Staphylococcus aureus*-activated astrocytes (36). In contrast, Arg1 expression was downregulated during *Salmonella* infection, and 15d-PGJ_2_ was unable to reverse this phenotype. iNOS and Arg1 compete for the same substrate, arginine, which is converted into nitric oxide or ornithine/urea, respectively. In general, iNOS is associated with inflammatory activation, whereas Arg1 is linked to anti-inflammatory responses and tissue repair, acting to counterbalance nitric oxide production (37). Overall, our results suggest that 15d-PGJ_2_ partially reprograms macrophage phenotypes during *Salmonella* infection, although it does not fully reverse their pro-inflammatory profile.

Since IL-1β release was one of the main targets modulated by 15d-PGJ_2_ in our studies, we further considered its role during *Salmonella* infection. Zigdon *et al*. demonstrated that depletion of IL-1β in *Salmonella*-infected mice led to a decrease in bacterial loads in the cecum and improved animal survival (38). The production of IL-1β is a hallmark of inflammasome activation (31). The role of inflammasome activation in the biology of *Salmonella* is controversial. Inflammasome activation, primarily through NLRC4, NLRP3, and non-canonical caspase-11 pathways, plays a central role in host defenses by promoting IL-1β and IL-18 secretion and inducing pyroptotic cell death. Indeed, *Salmonella* infection robustly activates NAIP/NLRC4 and NLRP3 inflammasomes through the recognition of flagellin and T3SS components, leading to caspase-1 activation and inflammatory cytokine release (39, 40). However, *Salmonella* has evolved sophisticated strategies to survive and replicate within host cells by tightly modulating innate immune responses, particularly those mediated by inflammasomes. Accumulating evidence indicates that *Salmonella* does not merely trigger the inflammasome pathway but instead establishes a finely tuned balance between inflammasome activation and evasion. For instance, bacterial factors such as FlgM regulate flagellin expression to limit NLRC4 detection, thereby enhancing virulence and promoting evasion of host protection (41). Consistent with this, persistent or unregulated flagellin expression leads to enhanced inflammasome activation and bacterial clearance, highlighting the importance of immune evasion for intracellular survival (42). Moreover, redundancy between the NLRC4 and NLRP3 pathways suggests that *Salmonella* encounters and modulates multiple inflammasome sensors during infection, allowing flexibility in host-pathogen interactions (39). Together, these findings support a model in which *Salmonella* dynamically modulates inflammasome activation - avoiding excessive pyroptosis that would eliminate its intracellular niche while still exploiting host inflammatory responses - thereby using inflammasome signaling not only as a target of immune evasion but as a process that can be strategically manipulated to favor its persistence and dissemination. Our results revealed that 15d-PGJ_2_ decreased the expression of inflammasome-related targets, including NLRP3 and GSDMD. Also, levels of active caspase-1 are reduced during treatment of *Salmonella*-infected cells with 15d-PGJ_2_. However, modulation of inflammasome activation alone, either by IL-1β or nigericin treatment, was not sufficient to reverse the effects of 15d-PGJ_2_ on bacterial colonization. Consistent with this, a previous study demonstrated that systemic *Salmonella* infection in caspase-1 or gasdermin D-deficient mice resulted in prolonged survival associated with reduced plasma levels of IL-1β, IL-6, and TNF-α (43). These results suggest that 15d-PGJ_2_ does not act through a single pathway but affects multiple inflammatory checkpoints in macrophages.

*Salmonella* is recognized by innate immune cells through pattern recognition receptors (PRRs) that detect PAMPs present in the bacteria cell. One of the main pathways described in *Salmonella* infection is the interaction between TLR4 and LPS, leading to activation of inflammatory signaling pathways, including NF-κB and the inflammasome (25, 27). During activation, TLR4 requires dimerization, which can occur either as a homodimer or through interaction with TLR2. Our data suggest that 15d-PGJ_2_ can partially downregulate the TLR4 pathway, including the expression of the receptor itself as well as TLR2, MyD88, p65, and NF-κB expression, when compared to untreated, infected cells. Supporting these findings, Li *et al*. showed that fisetin, a flavonoid, reduces intracellular *Salmonella* proliferation by inhibiting the TLR2/TLR4-NF-κB pathway in mice (44). In addition, exopolysaccharides derived from *Lactobacillus rhamnosus* GG were shown to suppress TLR4/NF-κB/MAPK signaling, leading to reduced intestinal injury in *Salmonella*-infected mice (45). Finally, probiotic treatment has been shown to reduce inflammation by inhibiting TLR4, NF-κB, and NLRP3 inflammasome signaling in porcine jejunal epithelial cells, resulting in protection of tight junction proteins such as ZO-1, occludin, and claudin-1 during *Salmonella* infection (46). These findings support the relevance of the TLR4-NF-κB-inflammasome axis during *Salmonella* infection. In our model, we observed that the combination of 15d-PGJ_2_ with a TLR4 antagonist further reduced *Salmonella* colonization and was associated with decreased IL-1β production and caspase-1 activation compared to 15d-PGJ_2_ treatment alone. Based on these results, we suggest that 15d-PGJ_2_ may act at the interface between TLR4 and inflammasome signaling, contributing to reduced bacterial colonization.

Although 15d-PGJ_2_ treatment downregulated several targets associated with pro-inflammatory responses, our results suggest that macrophages were not fully reprogrammed toward an anti-inflammatory phenotype. In addition, anti-inflammatory macrophage profiles have been associated with *Salmonella* persistence and resistance (47, 48). In contrast, pro-inflammatory responses are commonly linked to better control of bacterial burden, although frequently associated with increased tissue injury during infection (38, 43). Based on our findings, we believe that 15d-PGJ_2_ may act as a regulator that balances pro- and anti-inflammatory pathways, contributing to reduced *Salmonella* colonization. However, further studies are still necessary to better understand the exact mechanisms involved.

*In vivo*, 15d-PGJ_2_ also reduced *Salmonella* colonization, but only in the cecum. We believe this tissue specificity could be influenced by two factors. First, the cecum is the main site of *Salmonella*-induced inflammation during infection. Given the anti-inflammatory effects of 15d-PGJ_2_ extensively discussed herein, the cecum might represent the body site where the effects of 15d-PGJ_2_ may be more significant and therefore more easily observed. Second, it is important to note that 15d-PGJ_2_ was produced endogenously by both groups of animals in our experiments, including those that were not treated with this eicosanoid. Therefore, our mouse infection experiments, as executed, represent a comparison of *Salmonella* loads in two groups of animals that carried 15d-PGJ_2_. In one of the groups, only endogenously produced 15d-PGJ_2_ is present, whereas in the other group both endogenous and exogenous 15d-PGJ_2_ are present. In order to fully determine the impact of 15d-PGJ_2_ on *Salmonella* infection, animals deficient in the enzyme responsible for 15d-PGJ_2_ synthesis would need to be used.

Although the results of *Salmonella* burdens in mice described above fully support the notion that 15d-PGJ_2_ aids in the control of infection, the patterns of inflammatory gene expression *in vivo* were a bit less clear. In particular, we observed that the inflammatory gene expression in the cecum was different from what was observed in cultured macrophages. We observed increased expression of COX-2, TNF-α, TLR2, TLR4, MyD88, and NF-κB in infected mice treated with 15d-PGJ_2_, while GSDMD was decreased. These differences likely reflect the complexity of the intestinal environment, where multiple cell types and signaling inputs contribute to the final inflammatory response. In this context, 15d-PGJ_2_ does not appear to uniformly suppress inflammatory genes; rather, 15d-PGJ_2_ modulates the host response depending on the cell type and tissue context. Importantly, however, both *in vitro* and *in vivo* models show the same functional outcome: a reduction in *Salmonella* colonization. This suggests that 15d-PGJ_2_ regulates host-pathogen interactions through a multi-level mechanism, rather than through a single inflammatory pathway. Together, these results indicate that 15d-PGJ_2_ acts as a context-dependent immunomodulator that limits *Salmonella* colonization through the regulation of inflammatory pathways in cultured macrophages and *in vivo*.

## Acknowledgments

We thank Dr. Noraida Martinez-Rivera (Microscopy and Analytical Imaging Lab, University of Kansas) and Dr. Peter McDonald (Flow Cytometry Core, University of Kansas) for their technical support.

---

**Figure S1.**
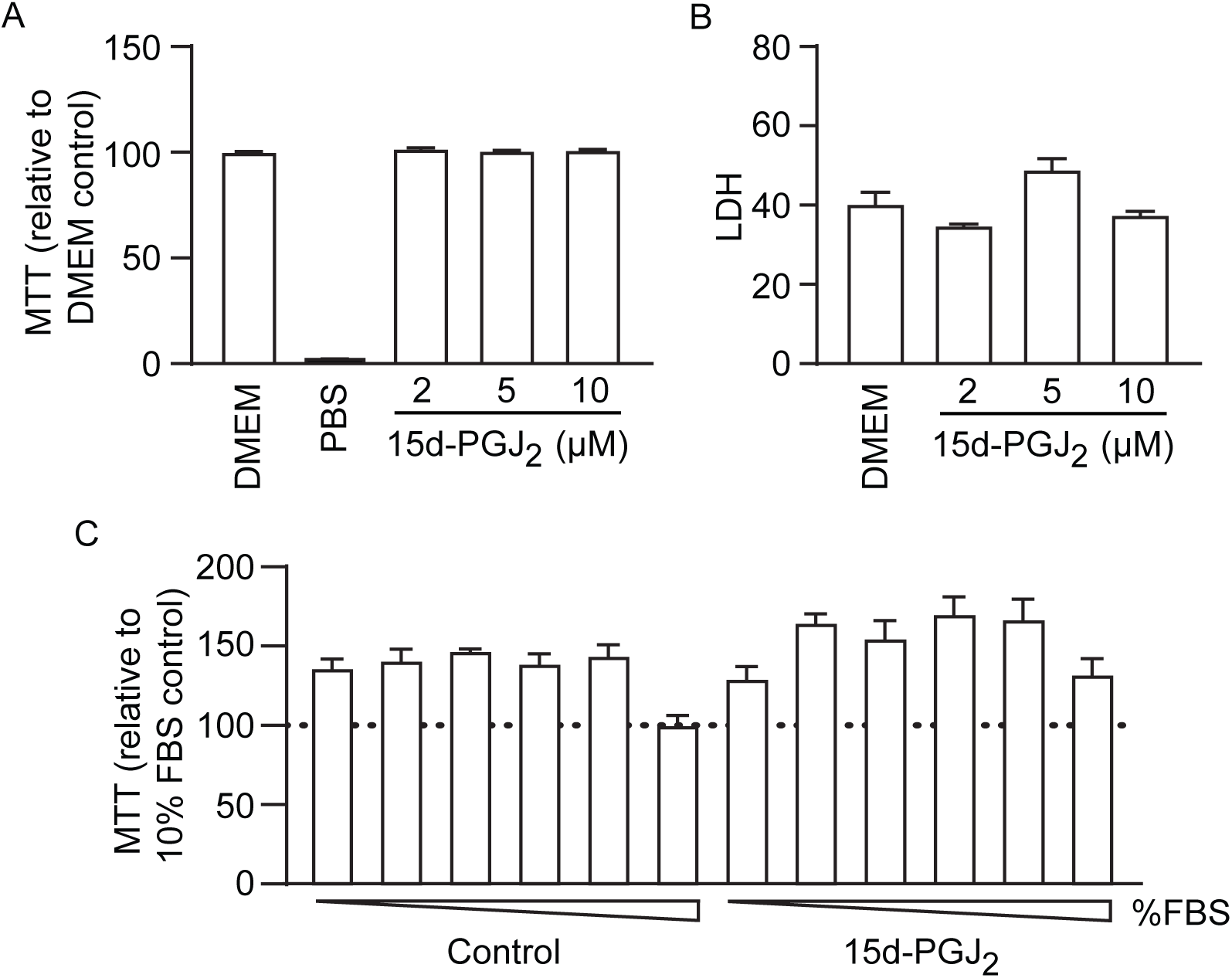
15d-PGJ_2_ does not affect macrophage viability. Metabolic (A) and cytotoxic (B) activities under different concentrations of 15d-PGJ_2_ were measured by MTT and LDH assays in macrophages after 24 hours of exposure, respectively. (C) Impact of 15d-PGJ_2_ (10 µM) treatment on metabolic activity over a range of FBS concentrations (0, 0.5, 1, 2, 5, and 10%), as measured by the MTT assay. The graphs plotted are representative of three independent experiments. Each bar represents the mean ± standard error of the mean of 5 replicates. Statistical analyses were performed using one-way ANOVA followed by the Tukey test. MTT, 3-(4,5-dimethylthiazol-2-yl)-2,5-diphenyltetrazolium bromide; LDH, lactate dehydrogenase; FBS, fetal bovine serum.

**Figure S2.**
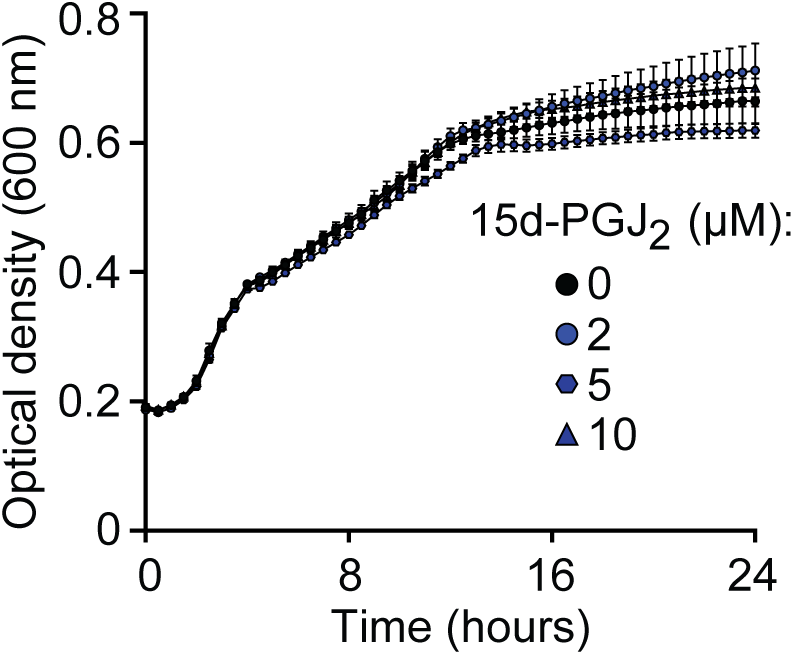
15d-PGJ_2_ does not affect *Salmonella* growth *in vitro*. (A) *Salmonella* was grown in LB broth supplemented with streptomycin (100 µg/mL) and different concentrations of 15d-PGJ_2_ at 37°C. Bacterial growth was monitored by measuring absorbance (OD600) over 24 hours using a microplate reader. The graph shows data from one representative experiment, selected from three independent experiments performed under the same conditions. Each independent experiment included 5 replicates per condition. Data points represent the means ± standard errors of the means (SEM) of the five replicates from the representative experiment shown. Statistical analyses were performed using the Student’s t-test.

**Figure S3.**
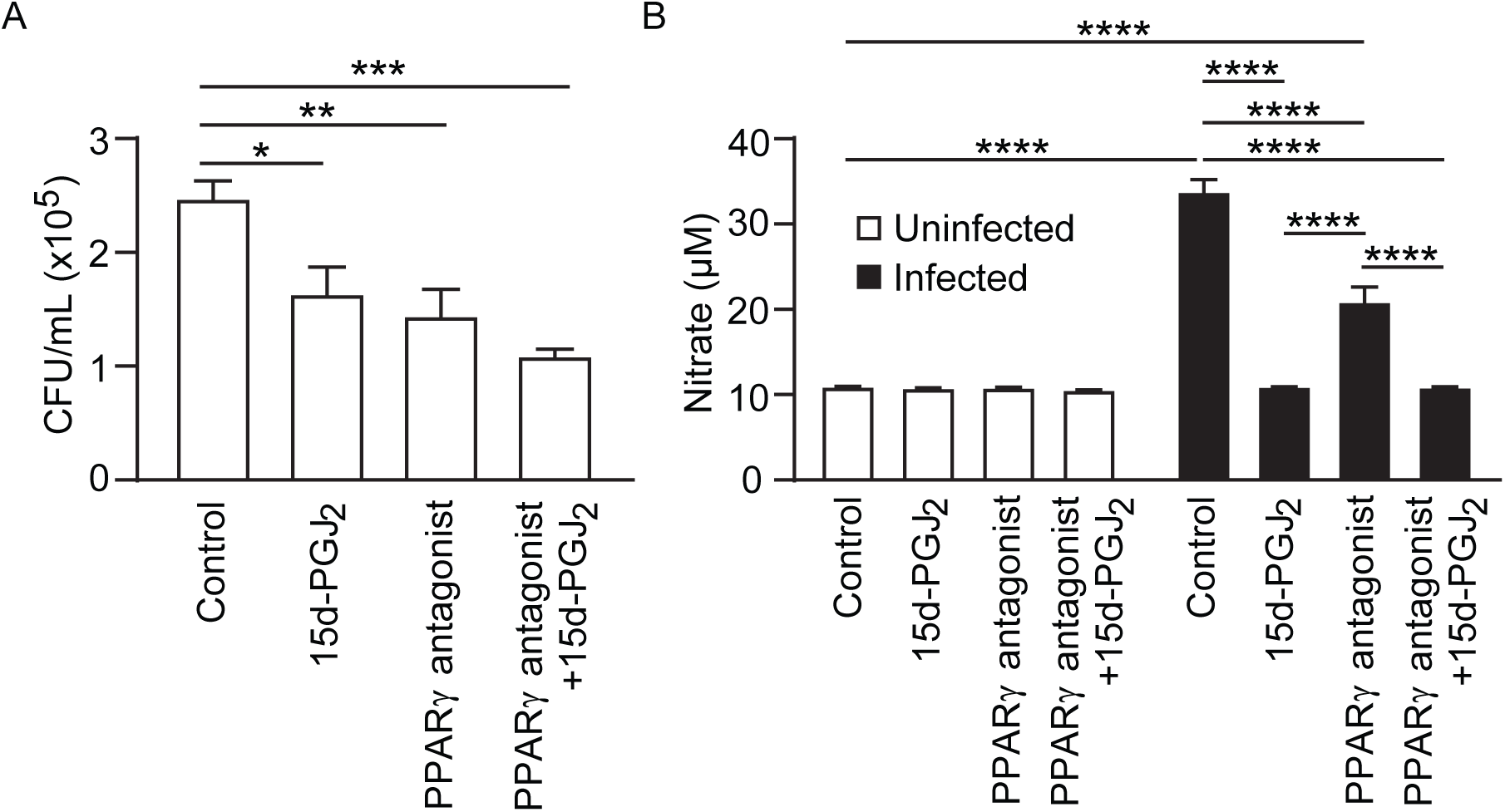
Effect of 15d-PGJ_2_ on Salmonella macrophage colonization is PPARγ independent. (A) The impact of the PPARγ antagonist T0070907 (1 µM) on macrophage colonization was evaluated by CFU quantification 24 hours after infection. (B) Nitrate levels were quantified using the Griess Reagent System Kit. An MOI of 10 was used. The graphs show data from one representative experiment out of two independent experiments performed under the same conditions. Each independent experiment included 5 replicates per condition. Bars represent the means ± standard errors of the means (SEM) of the five replicates from the representative experiment shown. Statistical analyses were performed using one-way ANOVA followed by the Tukey test. \**p*<0.05; \*\**p*<0.01; \*\*\**p*<0.001; \*\*\*\**p*<0.0001. H-PGDS, hematopoietic prostaglandin D synthase; CFU, colony-forming units; IL, interleukin; n.d., not detected.

**Figure S4.**
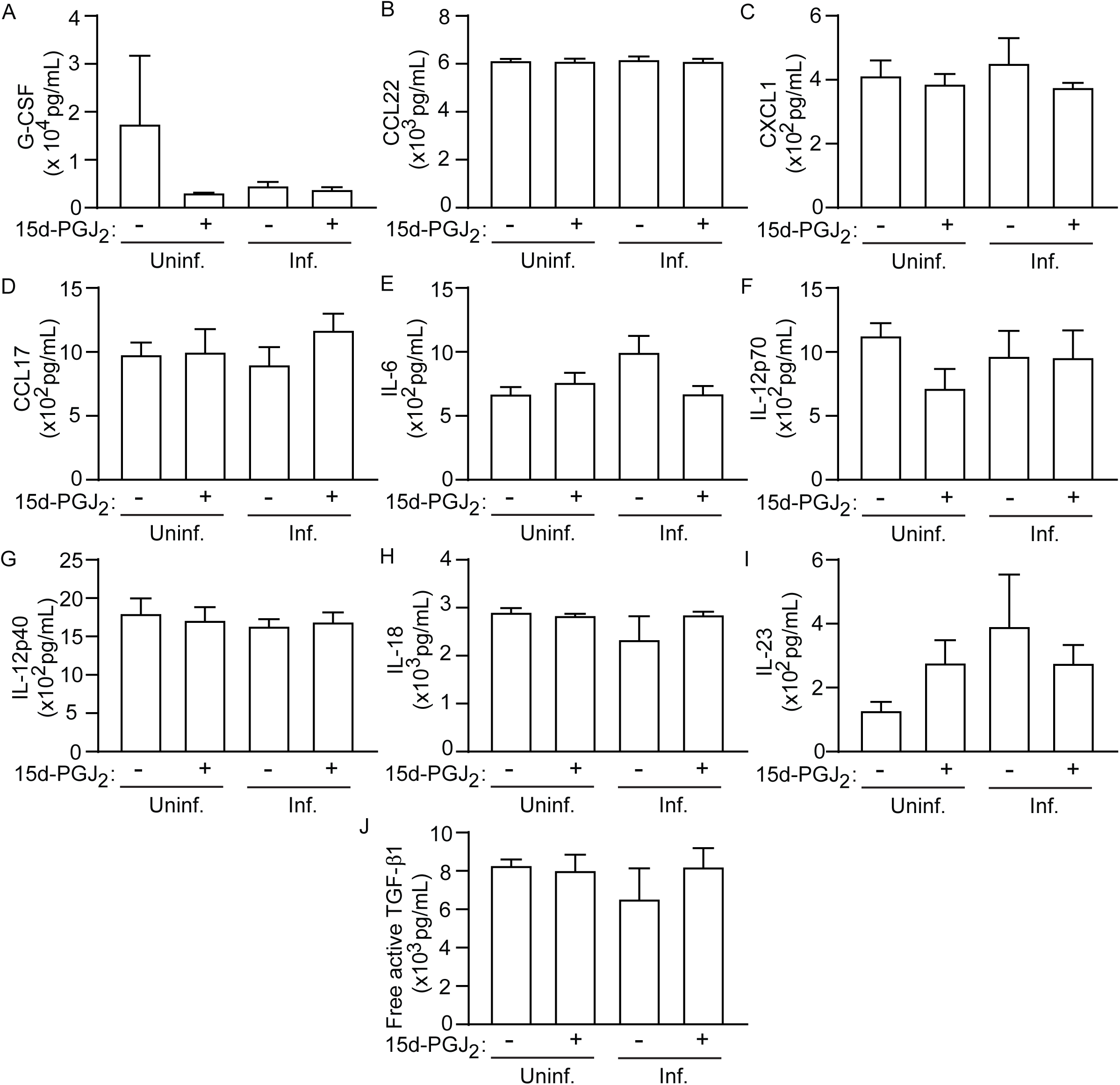
15d-PGJ_2_ has no significant impact on the production of selected cytokines and chemokines in *Salmonella*-infected macrophages. The effect of 15d-PGJ_2_ (10 µM) on (A) G-CSF, (B) CCL22, (C) CXCL1, (D) CCL17, (E) IL-16, (F) IL-12p70, (G) IL-12p40, (H) IL-18, (I) IL-23, and (J) free active TGF-β1 was analyzed 24 hours after infection by LEGENDplex™ Mouse Macrophage/Microglia Panel Kit. An MOI of 10 was used. Data shown are from a single experiment performed with 4 replicates per condition. Bars represent the means ± standard errors of the means (SEM) of the four replicates. Statistical analyses were performed using one-way ANOVA followed by the Tukey test. G-CSF, granulocyte colony-stimulating factor; CCL22, C-C motif chemokine ligand 22; CXCL1, chemokine (C-X-C motif) ligand 1; CCL17, C-C motif chemokine ligand 17; IL, interleukin; TGF-β1, transforming growth factor beta 1.

**Figure S5.**
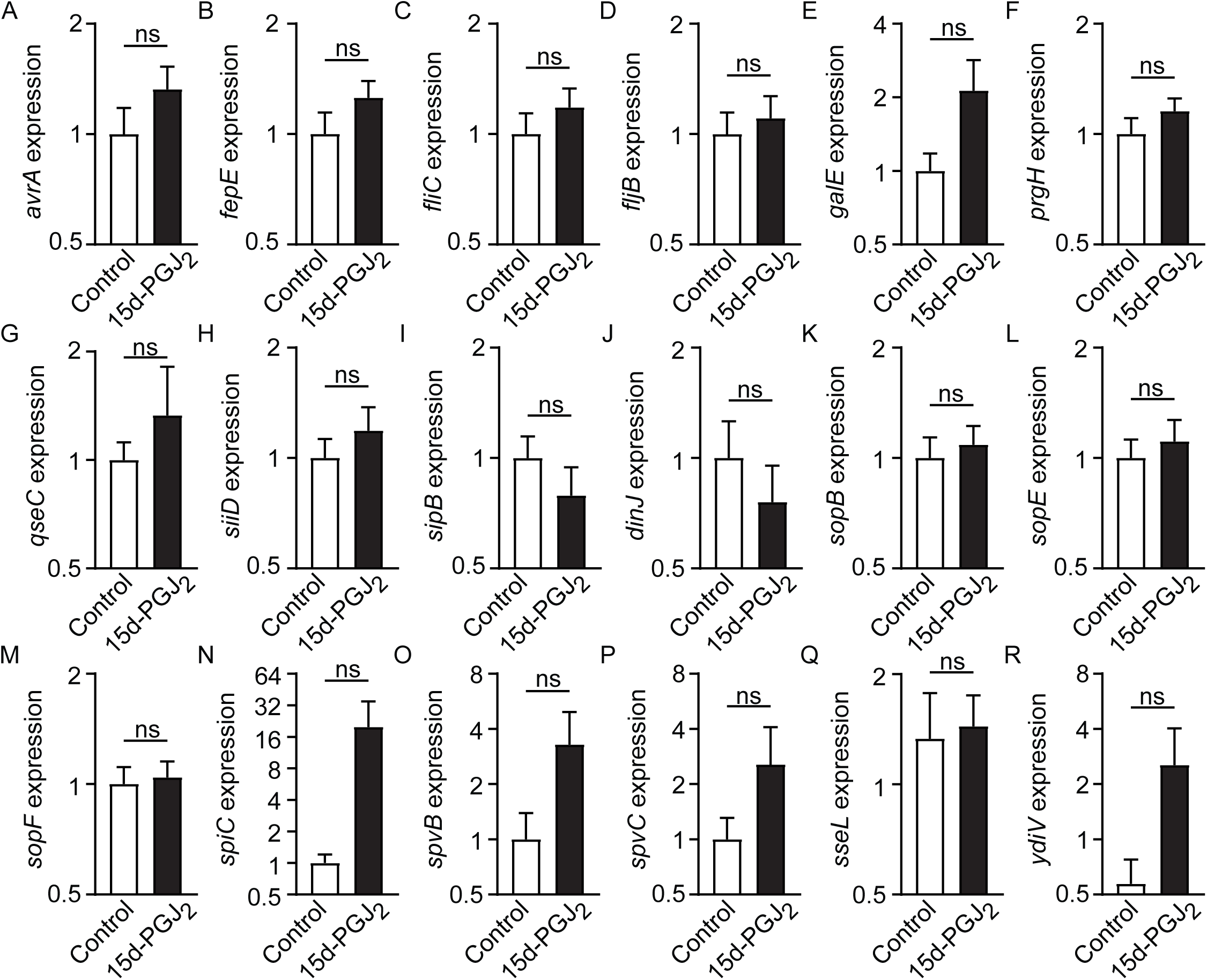
Effect of 15d-PGJ_2_ treatment on the expression of inflammasome-related *Salmonella* genes. The effect of 15d-PGJ_2_ (10 µM) on (A) *avrA*, (B) *fepE*, (C) *filC*, (D) *fijB*, (E) *galE*, (F) *prgH*, (G) *qseC*, (H) *siiD*, (I) *sipB*, (J) *dinJ*, (K) *sopB*, (L) *sopE*, (M) *sopF*, (N) *sipC*, (O) *spvB*, (P) *spvC*, (Q) *sseL*, and (R) *ydiV* expression was analyzed by qRT-PCR in *Salmonella*. *Salmonella* cultures were grown in LB broth supplemented with streptomycin (100 µg/mL) overnight. Subcultures (1:200) were supplemented with 15d-PGJ_2_ (10 µM) and incubated at 37°C until the mid-log phase of growth was reached (4 hours), prior to qRT-PCR analysis. Data shown are pooled from two independent experiments, each performed with 4 replicates per condition. Bars represent the means ± standard errors of the means (SEM) of the pooled data. Statistical analyses were performed using the Student’s t-test. ns, non-significant.

**Figure S6.**
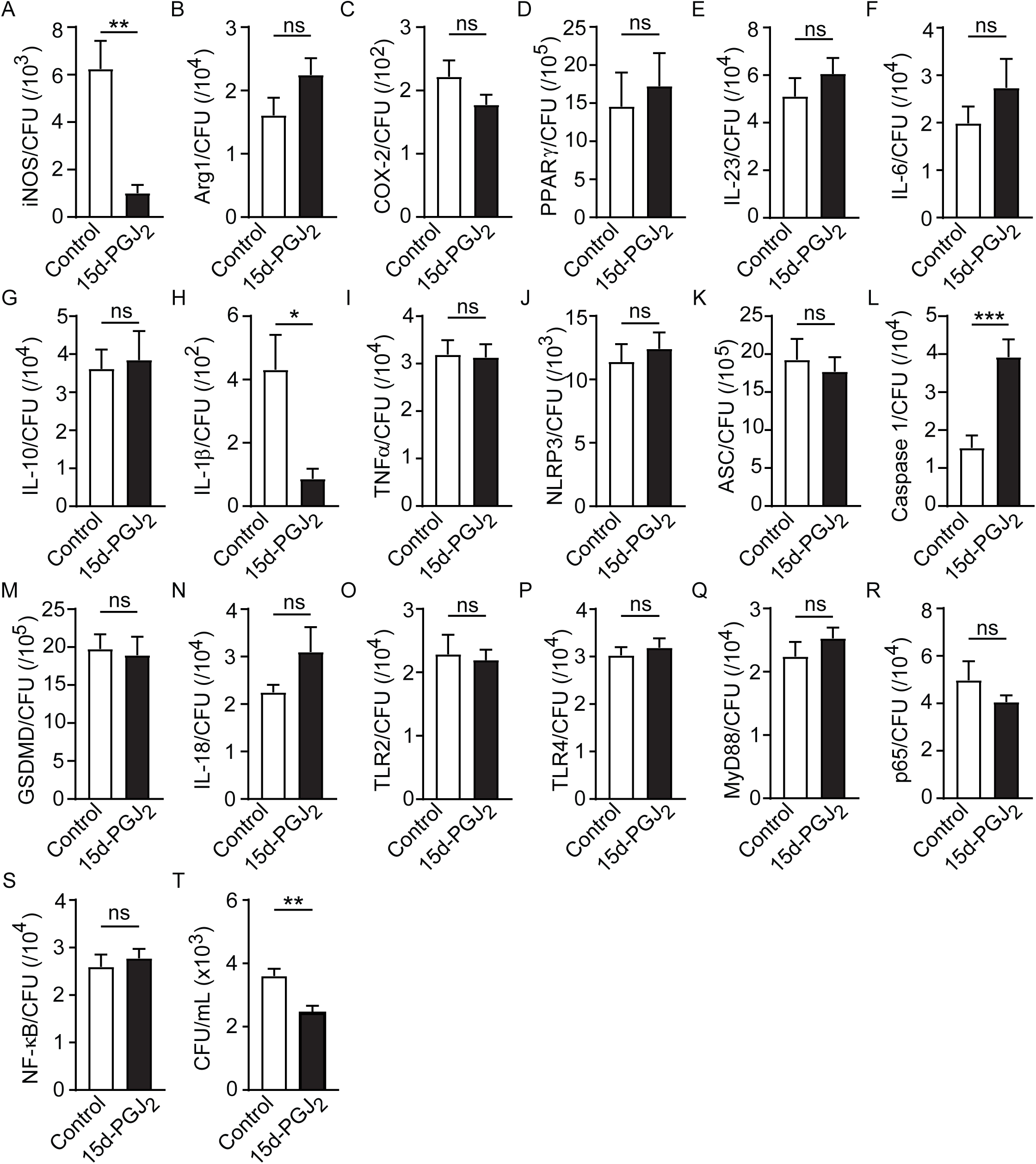
Effect of 15d-PGJ_2_ treatment on the expression of inflammatory-related genes in macrophages after normalization by bacterial loads. Expression levels of (A) iNOS, (B) Arg1, (C) COX-2, (D) PPARγ, (E) IL-23, (F) IL-6, (G) IL-10, (H) IL-1β, (I) TNF-α, (J) NLRP3, (K) ASC, (L) caspase-1, (M) GSDMD, (N) IL-18, (O) TLR2, (P) TLR4, (Q) MyD88, (R) p65, and (S) NF-κB were measured and normalized to average bacterial counts (CFU). (T) Effect of 15d-PGJ_2_ (10 µM) on *Salmonella* colonization of macrophages 24 hours after infection. An MOI of 10 was used. Data shown are pooled from two independent experiments, each performed with 5 replicates. Bars represent the means ± standard errors of the means (SEM) of the pooled data. Statistical analyses were performed using the Student’s t-test. ns, non-significant; \**p*<0.05; \*\**p*<0.01; \*\*\**p*<0.001. iNOS, inducible nitric oxide synthase; Arg1, arginase 1; COX-2, cyclooxygenase-2; PPARγ, peroxisome proliferator-activated receptor gamma; IL, interleukin; TNF-α, tumor necrosis factor alpha; NLRP3, NOD-like receptor family pyrin domain-containing 3; ASC, apoptosis-associated speck-like protein containing a CARD; GSDMD, gasdermin D; TLR, Toll-like receptor; MyD88, myeloid differentiation primary response 88; NF-κB, nuclear factor kappa-light-chain-enhancer of activated B cells; CFU, colony-forming units.

**Figure S7.**
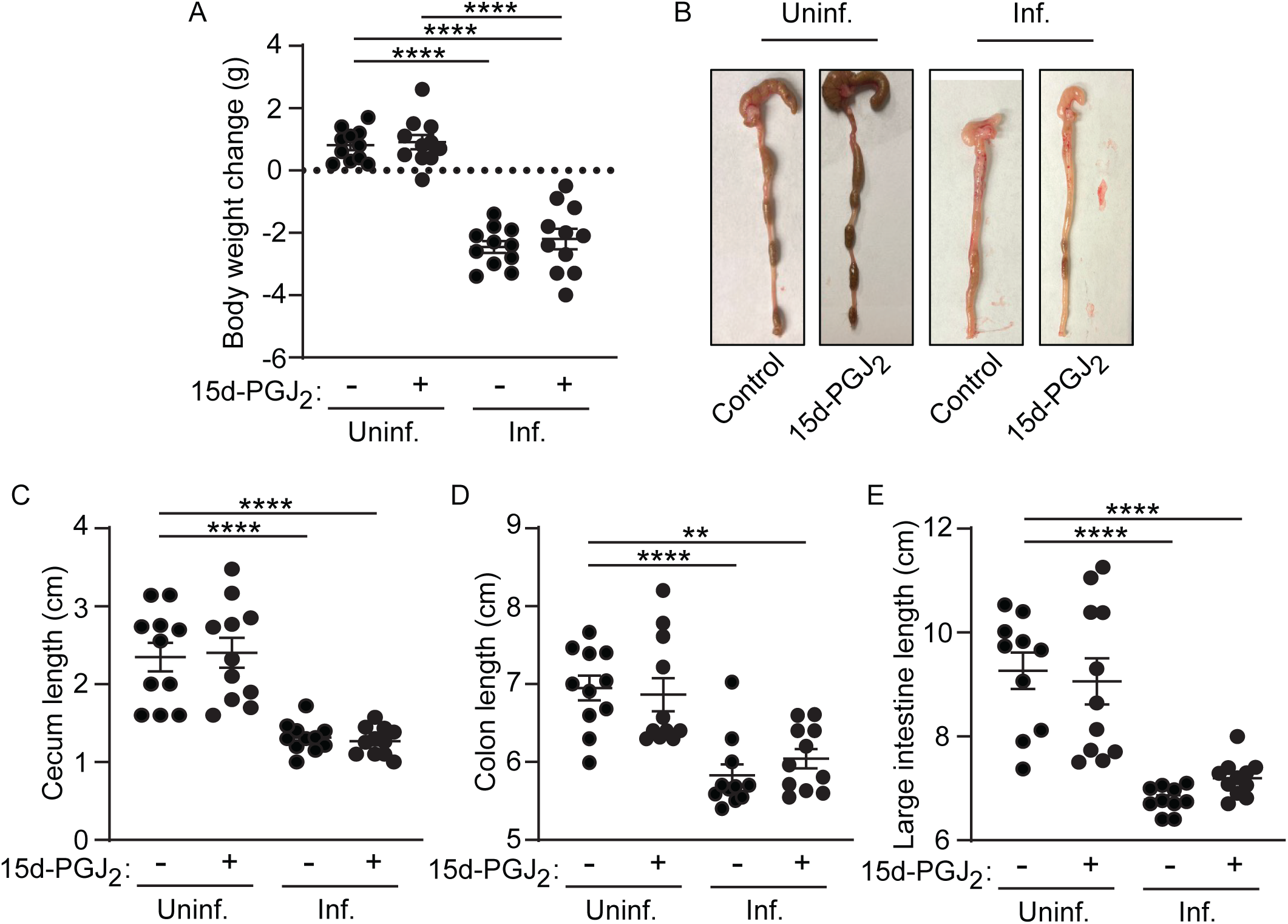
15d-PGJ_2_ treatment does not alter body weight or intestinal length in *Salmonella*-infected mice. (A) Ratios of final to initial body weight of mice. (B) Representative images of the large intestines. The graphs represent the lengths of the cecum (C), colon (D), and large intestine (E), measured in centimeters. The 15d-PGJ_2_ dose used was 1 mg/kg. Dots represent individual animals, and the data shown are pooled from two independent experiments, with 5 mice per experiment. Outliers were identified and removed using the ROUT method (Q = 1%) prior to statistical analysis. Statistical analyses were performed using one-way ANOVA followed by the Tukey test. \*\**p*<0.01; \*\*\*\**p*<0.0001.

