## Supplemental Table 1 for "15-deoxy-Δ^12,14^-prostaglandin J_2_ limits *Salmonella* infection through regulation of host TLR4 signaling and inflammasome activation"

**Table S1: Sequences of primers used in the study.**

| Target | Sequence (5’-3’) |
| --- | --- |
| *Eukaryotic primers* |  |
| TNF-α | Forward TCTCCTTCCTGATCGTGGCA  Reverse CAGCTTGAGGGTTTGCTACAAC |
| IL-6 | Forward CATCCTCGACGGCATCTCAG  Reverse TCACCAGGCAAGTCTCCTCA |
| IL-10 | Forward GTAGAAGTGATGCCCCAGGC  Reverse CACCTTGGTCTTGGAGCTTATT |
| Arg-1 | Forward TTTCTCAAAAGGACAGCCTCG  Reverse ACAGACCGTGGGTTCTTCAC |
| PPARγ | Forward CTCTGTTTTATGCTGTTATGGGTGA  Reverse GGTCAACAGGAGAATCTCCCAG |
| iNOS | Forward GGAATCTTGGAGCGAGTTGT  Reverse GCAGCCTCTTGTCTTTGACC |
| Myd88 | Forward GCCAGAGTGGAAAGCAGTGT  Reverse TATCGTTGGGGCAGTAGCAG |
| p65 | Forward TAACAGCAGGACCCAAGGAC  Reverse AGCCCCTAATACACGCCTCT |
| NF-κB | Forward GCCAGACACAGATGATCGCC  Reverse GTTTCGGGTAGGCACAGCAA |
| COX-2 | Forward TTCCAATCCATGTCAAAACCGT  Reverse AGTCCGGGTACAGTCACACTT |
| TLR4 | Forward CCTGATGACATTCCTTCT  Reverse AGCCACCAGATTCTCTAA |
| IL-1β | Forward TGGACCTTCCAGGATGAGGACA  Reverse GTTCATCTCGGAGCCTGTAGTG |
| TLR2 | Forward ACCAAGATCCAGAAGAGCCA  Reverse CATCACCGGTCAGAAAACAA |
| Asc | Forward GTGTTTACTCTCTGGGATGTTTTTG  Reverse GTCTGTGGAATTTAGGTGTTGGA |
| Nlrp3 | Forward GGGAGACCGTGAGGAAAGGA  Reverse CCAAAGAGGAATCGGACAACAAA |
| IL-18 | Forward TGCAGACTGGTTGGCATCAA  Reverse CTGATGCTGGAGGTTGCAGA |
| GSDMD | Forward TGGTGAAGCACGTCTTGGAA  Reverse GTGGGGATCAGAGACGTTGG |
| Caspase 1 | Forward TGCAGACTGGTTGGCATCAA  Reverse CTGATGCTGGAGGTTGCAGA |
| IL-23 | Forward ACATGCACCAGCGGGACATA  Reverse CTTTGAAGATGTCAGAGTCAAGCAG |
| *Bacterial primers* |  |
| *avrA* | Forward TCGAGCTGGACATTCAACGAAG  Reverse GTTCTTCACCACACAGACGTTCAC |
| *fepE* | Forward TGCTTCTGTCCTTTCTGCTGC  Reverse CGATCAACGCTTACCTCCATATCC |
| *fliC* | Forward TCAACGGCGTGAAAGTCCTG  Reverse GCACATTCAGCGTATCCAGAC |
| *fljB* | Forward CGCCAACGACGGTGAAACTATC  Reverse ACCACCCGTAGCCGCTTTAATAG |
| *galE* | Forward ATGATTACCCGACCGAGGATGG  Reverse AAAGGCGTTGACCACATCCAG |
| *prgH* | Forward AAGCGTATCTCTATCTGGCTGG  Reverse CTGCTGCGGTAACATCGTCCAT |
| *qseC* | Forward GCTATCTGAAGGATGACAACGACC  Reverse ATGAAGGGAAGGGCGATCAGC |
| *siiD* | Forward AGCCGTTAAACTGGATGTGC  Reverse TGGCGTCAACAGTCATACCTGG |
| *sipB* | Forward TGGGACTTGCGGTAATGGTG  Reverse CGCTTTGGTAATCGCCTTGC |
| *sopB* | Forward GCGGCAGTAGCGTTTAGAGATG  Reverse CATACGCCCTTTCCCTCATAAGCAC |
| *sopE* | Forward GCTCATCAGCAAAGGGATCAAC  Reverse GGATGCCTGCTGATGTTGATTG |
| *sopF* | Forward CGTGAGTGTGGTGCCGAAATAG  Reverse GGGGCATAAGGTGAATCTG |
| *spiC* | Forward CGAAGGTAATAGCCGATCCTGG  Reverse GCAAGCAGTAGTGTCACATAGG |
| *spvB* | Forward CGGAGAAGTGCTGGTTCAAACG  Reverse CCAGAAATCATCGCCGTTGC |
| *spvC* | Forward GCTGGATGTGCCTGACTATTC  Reverse CACTGATGTGGAACTTGTCACC |
